# Long Non-Coding RNA *LncKdm2b* Regulates Cortical Neuronal Differentiation by *Cis*-Activating *Kdm2b*

**DOI:** 10.1101/459289

**Authors:** Wei Li, Wenchen Shen, Bo Zhang, Kuan Tian, Yamu Li, Lili Mu, Zhiyuan Luo, Xiaoling Zhong, Xudong Wu, Ying Liu, Yan Zhou

**Affiliations:** College of Life Sciences, Medical Research Institute at School of Medicine, Wuhan University, Wuhan 430072, China; Department of Cell Biology, Tianjin Medical University, Qixiangtai Road 22, Tianjin 300070, China

## Abstract

The mechanisms underlying spatial and temporal control of cortical neurogenesis of the brain are largely elusive. Long non-coding RNAs (lncRNAs) have emerged as essential cell fate regulators. Here we found *LncKdm2b* (also known as *Kancr*), a lncRNA divergently transcribed from a bidirectional promoter of *Kdm2b*, is transiently expressed during early differentiation of cortical projection neurons. Interestingly, *Kdm2b’s* transcription is positively regulated *in cis* by *LncKdm2b*, which has intrinsic-activating function and facilitates a permissive chromatin environment at the *Kdm2b’s* promoter by associating with hnRNPAB. Lineage tracing experiments and phenotypic analyses indicated *LncKdm2b* and *Kdm2b* are crucial in proper differentiation and migration of cortical projection neurons. Moreover, KDM2B exerts its role relying on its leucine-rich repeats (LRR) but independent of its PRC1-related function. These observations unveiled a lncRNA-dependent machinery in regulating cortical neuronal differentiation.

## Introduction

The mammalian cerebral cortex, also known as the neocortex, is a six-layered structure and responsible for performing the most sophisticated cognitive and perceptual functions such as sensory perception, generation of motor commands, conscious thought and language. The adult neocortex comprises a plethora of projection neurons, interneurons and glial cells. Projection neurons (PNs) are the main functional units, expressing excitatory neurotransmitters, with their long axons projecting into subcortical regions or contralateral cortex of the brain. In mice, cortical PNs are largely generated between embryonic (E) day 11.5 to E17.5 indirectly from radial glial progenitor cells (RGPCs), whose nuclei lie in the region close to the lateral ventricles, ventricular zone (VZ). RGPCs usually divide asymmetrically to self-renew and simultaneously give rise to intermediate progenitor cells (IPCs), which are multipolar and reside basally to RGPCs in the subventricular zone (SVZ). IPCs divide symmetrically to generate either two IPCs or two postmitotic PNs. PNs then migrate radially along the basal processes of RGPCs to propagate the cortical plate (CP) in the basal part of the cortex, which eventually forms cortical layers (Fietz and Huttner, 2011; Kwan et al., 2012). Many cellular and molecular aspects governing cortical neurogenesis have been extensively studied, including cell-autonomous and non-autonomous regulation of RGPCs’ asymmetric cell division, neuronal fate commitment, as well as PNs’ radial migration (Ayala et al., 2007; Greig et al., 2013; Imayoshi and Kageyama, 2014). However, mechanisms that control the initial numbers and proliferation rates of RGPCs, as well as the proliferative or neurogenic choices of IPCs, are largely elusive (Greig et al., 2013; Homem et al., 2015).

Recent studies indicate a few long non-coding RNAs could be essential cell fate regulators in development (Grote et al., 2013; Klattenhoff et al., 2013). Long non-coding RNAs (lncRNAs), defined as RNAs longer than 200 nucleotides but lacking protein-coding potentials, are abundant in brain and display cell-type-, and developmental stage-specific expression patterns compared to protein-coding transcripts (Aprea et al., 2013; Belgard et al., 2011; Mercer et al., 2010; Molyneaux et al., 2015). LncRNAs may regulate gene transcription by recruiting transcription factors, RNA-binding proteins and chromatin-remodeling machineries to the site of transcription and creating a locus-specific environment (Lin et al., 2014; Ng et al., 2013; Wang et al., 2015). LncRNAs are often derived from bidirectional promoters, such that initiating Pol II can generate divergently-oriented transcripts simultaneously, the sense (protein-coding mRNA) direction or the upstream-antisense (divergent non-coding) direction, with these mRNA/divergent lncRNA pairs having coordinated expression (Lepoivre et al., 2013; Scruggs and Adelman, 2015; Sigova et al., 2013). Moreover, the transcription of divergent lncRNAs could affect the expression of their neighboring protein-coding transcripts *in cis* (Luo et al., 2016; ørom et al., 2010). Anti-sense promoters could serve as platforms for transcription factor (TF) binding and facilitate establishment of proper chromatin architecture to regulate sense-strand mRNA expression (Scruggs and Adelman, 2015; Scruggs et al., 2015). Although divergent lncRNAs are prevalent in both embryonic and adult nervous system, only a few functional divergent lncRNAs have been characterized, including roles of *Emx2OS* and in regulating the expressions of their neighboring protein-coding transcripts *Emx2*, an essential cortical RGPC gene (Noonan et al., 2003; Spigoni et al., 2010). Furthermore, these are largely in *vitro* studies and it’s still lack of in *vivo* evidence showing the significance of divergent lncRNAs in cortical neuronal differentiation (Wang et al., 2017).

Here we characterized *LncKdm2b* (also known as *Kancr - **<u>K</u>**dm2b* upstream-**<u>a</u>**ntisense **<u>n</u>**on-**<u>c</u>**oding **<u>R</u>**NA), a divergent lncRNA that can positively regulate the transcription of *Kdm2b in cis*. Both *LncKdm2b* and *Kdm2b* are transiently expressed in committed neuronal precursors and newborn cortical PNs and essential for their proper differentiation. *LncKdm2b cis*-regulates *Kdm2b’s* expression and facilitates a permissive chromatin environment by binding to hnRNPAB. Our findings advance understandings of molecular events that govern cortical neuronal differentiation and might have general implications in regulation of cell differentiation.

## Results

### *LncKdm2b* and *Kdm2b* are Transiently Expressed in Committed Neuronal Precursors and Newborn Cortical Projection Neurons

In an effort to identify pairs of divergent lncRNA/protein-coding transcript that exert roles in cortical neurogenesis of the mouse brain, we analyzed a database comprising both in-house and publicized transcriptome data of developing mouse cerebral cortex (dorsal forebrain). In-house data are RNA-seq data from embryonic (E) day 10.5 and E12.5 dorsal forebrain. We also included RNA-seq data of mouse embryonic stem cells (mESCs), mESCs derived neural progenitor cells (NPCs), and tissues from later stages of cortical development including E14.5 ventricular zone (VZ), subventricular and intermediate zone (SVZ/IZ) and cortical plate (CP), E17.5 and adult cortex (Ayoub et al., 2011; Dillman et al., 2013; Guttman et al., 2010; Ramos et al., 2013). Interestingly, protein-coding genes associated with divergent lncRNAs within 5 kilobase from their transcription start sites (TSS) are highly enriched for signatures including transcription, cell cycle progression and catabolic process (Figure 1 - figure supplement 1A, Supplementary file 1 - Table 1), indicating their related roles (Ponjavic et al., 2009). One of these pairs is *Kdm2b* and its divergent non-coding transcript *LncKdm2b* (also known as *Kancr* and *A930024E05Rik)* (Diez-Roux et al., 2011; Liu et al., 2017; Saba et al., 2015). *LncKdm2b* is transcribed at 262 base pair upstream of *Kdm2b’s* TSS, and is predicted to be a lncRNA according to its low score in coding potential and inability to translate proteins (Figure 1 - figure supplement 1B-1C). The expression of *LncKdm2b* peaks in E14.5 SVZ/IZ, where IPCs and migrating PNs reside. Similarly, the expression of *Kdm2b* in E14.5 SVZ/IZ is slightly higher than that in E14.5 VZ and CP (Figure 1 - figure supplement 1D). Notably, *LncKdm2b* is expressed at higher levels than *Kdm2b* in E14.5 VZ and SVZ/IZ and at comparable levels in other stages (Figure 1 - figure supplement 1D), which is contradictory to the common notion that divergent lncRNAs are expressed at much lower levels than their neighboring protein-coding transcripts (Sigova et al., 2013). Consistently, quantitative RT-PCR and immunoblotting experiments showed expression levels of both KDM2B and *LncKdm2b* peak in E12.5 and E14.5 dorsal forebrains, with much lower levels in E10.5 and adult stages (Figure 1 - figure supplement 1E-1F, Figure 1 - figure supplement 1M). This pattern is quite similar to those of *Tbr2, Dcx, Unc5d* and *Neurod1*, markers for IPCs and immature PNs (Figure 1 - figure supplement 1G-1M). Northern blot detected a ~1.8 kb band in poly(A) RNAs extracted from E14.5 and E16.5 cortices (Figure 1 - figure supplement 1N). Through analyzing the ENCODE database (Yue et al., 2014), we found the genomic region spanning the promoter of *Kdm2b* and its immediate upstream region that transcribes *LncKdm2b* is evolutionarily conserved across mammals, and is associated with Pol II (RNA polymerase II) and H3K4me3 in E14.5 mouse brain, indicating active transcription at this condition (Figure 1A). *In situ* hybridization (ISH) revealed that both *LncKdm2b* and *Kdm2b* are predominantly expressed in the upper SVZ of the E16.5 dorsal forebrain, with the apical side of ISH signals overlapping with TBR2, an SVZ marker labeling intermediate cortical neural precursors (IPCs) (Figure 1B, S1O-S1P); and basal side overlapping with TUJ1, a marker for fate-determined pyramidal neurons (Figure 1 - figure supplement 1P). These data suggest both *LncKdm2b* and *Kdm2b* are transiently expressed in committed IPCs and freshly differentiated projection neurons during the peak of cortical neurogenesis.

**Figure 1.**
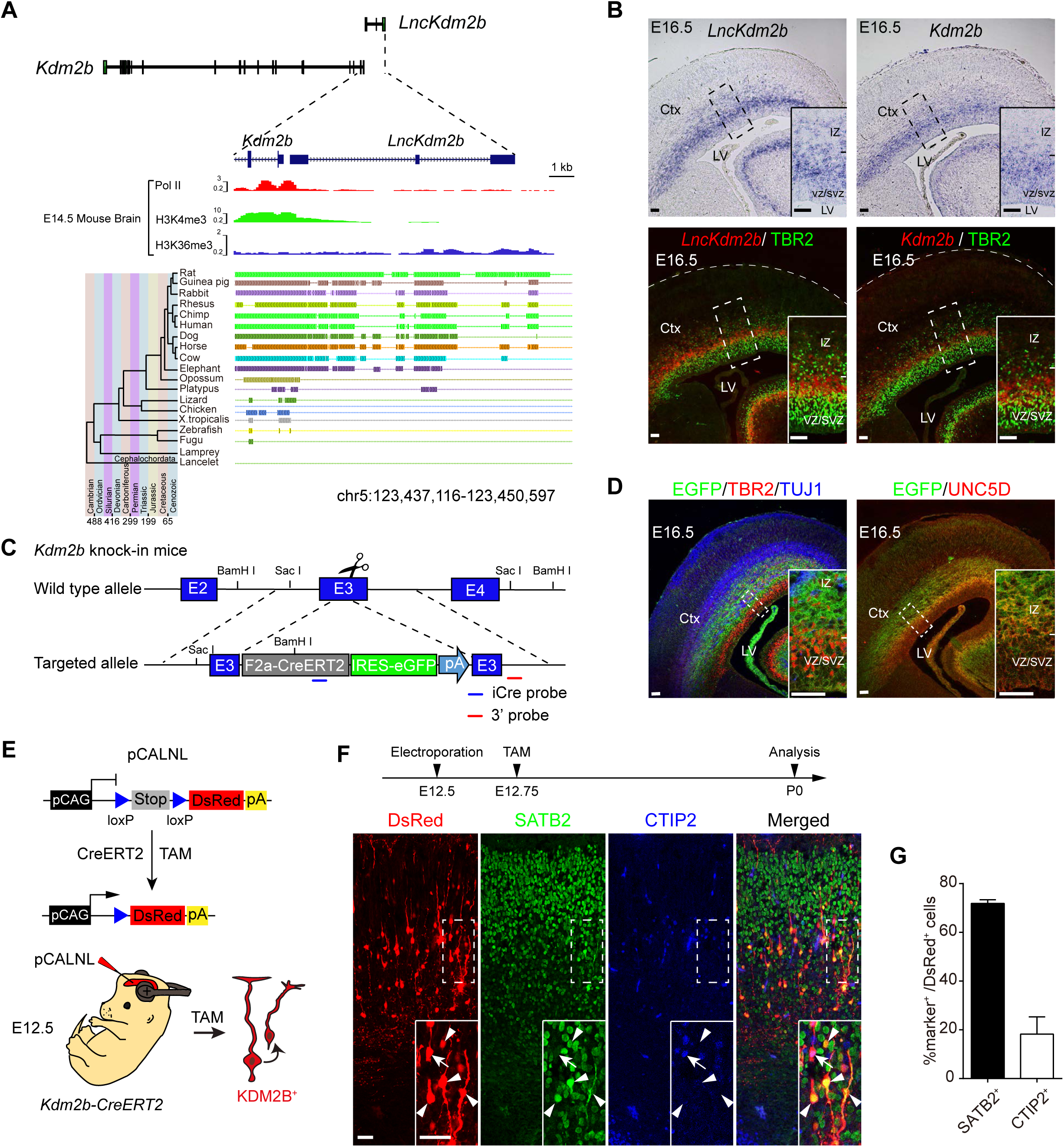
*LncKdm2b* and *Kdm2b* are Transiently Expressed in the Developing Mouse Embryonic Cortex. (A) Schematic illustration of the mouse *LncKdm2b/Kdm2b* locus. The top tracks depict ChIP-seq signals for Pol II, H3K4me3 and H3K36me3 in E14.5 mouse brain. Bottom tracks depict a parallel genomic alignment of 19 vertebrates to the mouse genome (UCSC mm9) at the *LncKdm2b* locus. Shaded lines indicate conserved sequences. (B) Top: *In situ* hybridization (ISH) of *LncKdm2b* (left) and *Kdm2b* (right) on coronal sections of E16.5 mouse dorsal forebrains. Bottom: Immunofluorescent staining for TBR2 (green) on ISH sections of *LncKdm2b* (left, red) and *Kdm2b* (right, red) on coronal sections of E16.5 mouse dorsal forebrains. (C) A schematic diagram illustrates the strategy for generating *Kdm2b^CreERT2^* knock-in mice line. (D) Left: Immunofluorescent staining for EGFP (green), TBR2 (red), and TUJ1 (blue) on cortical sections of E16.5 heterozygous *Kdm2b^CreERT2^* knock-in mice. Right: Immunofluorescent stainings for EGFP (green) and UNC5D (red) on cortical sections of E16.5 heterozygous *Kdm2b^CreERT2^* knock-in mice. (E) A schematic diagram illustrates the strategy for lineage tracing of Kdm2b-expressing cortical cells using *in utero* electroporation. (F) E12.5 *Kdm2b^CreERT2/+^* knock-in cortices were electroporated with conditional DsRed-expressing plasmids (pCALNL), followed by tamoxifen (TAM) injection at E12.75 and analyses for SATB2 (green) and CTIP2 (blue) expression at P0. Arrowheads indicate DsRed+, SATB2+ cells. Arrows indicate DsRed+, CTIP2+ cells. (G) Quantification of SATB2 or CTIP2 expression in DsRed+ recombined cells (F). A total of 711 cells from 3 animals were analyzed. Data shown are the mean + SD.Scale bars, 50 μm. Boxed areas are enlarged at the bottom-right corners in (B), (D) and (F). Ctx, cortex; LV, lateral ventricle; VZ, ventricular zone; SVZ, subventricular zone; IZ, intermediate zone. pA, polyA. See also Figure 1 - figure supplement 1 and 2.

### Kdm2b-expressing Cortical Cells are Fated to be Cortical Projection Neurons

To further validate *Kdm2b’s* expression and the fate of *Kdm2b*-expressing cells during cortical neurogenesis, we generated a knock-in mouse line, *Kdm2b-F2a-CreERT2-IRES-EGFP* (referred to *Kdm2b^CreERT2^*), in which the *F2a-CreERT2-IRES-EGFP* cassette was inserted in frame into the third exon of *Kdm2b* (Figure 1C). Southern blotting and genomic PCR validated the predicted genomic modification (Figure 1 - figure supplement 1Q). Expressions of CreERT2 and EGFP are driven by the endogenous *Kdm2b* promoter, which would allow us to perform detailed expression analyses and lineage tracing experiments for *Kdm2b*. Brain sections from embryos derived from mating of *Kdm2b^CreERT2/+^* with wild-type (WT) C57/B6 were subjected to immunofluorescent staining. Consistent with ISH experiments, EGFP+ cells reside in upper SVZ and lower intermediate zone (IZ), overlapping with both TBR2+ IPCs and TUJ1+ projection neurons (Figure 1D, S1R). Moreover, a large portion of EGFP+ cells also overlap with UNC5D, a marker for multipolar cells in embryonic SVZ/IZ and layer IV projection neurons (Figure 1 - figure supplement 1S). Notably, EGFP+ signals extend more basally than *Kdm2b* or *LncKdm2b* ISH signaling, probably because EGFP protein is more stable than transcripts of *Kdm2b* or *LncKdm2b*. We next bred *Kdm2b^CreERT2/+^* with the *Ai14 (Rosa-CAG-LoxP-STOP-LoxP-tdTomato-WPRE)* reporter mice. Pregnant female mice were injected with tamoxifen at various stages to enable the excision of the STOP cassette, thus leading to tdTomato expression in the progenies of *Kdm2b*-expressing cells. Cortices were collected from E16.5 and newborn (P0) pups for immunofluorescent staining of SATB2 (a marker for layer 2-4 callosal neurons) and CTIP2 (a marker for layer 5 subcortical neurons). Interestingly, most tdTomato-positive cells express either SATB2 (51.0 ± 2.5% at E16.5, 63.1 ± 2.5% at P0) or CTIP2 (20.7 ± 5.4% at E16.5, 7.0 ± 2.3% at P0), suggesting the progenies of *Kdm2b*-expressing cells are largely projection neurons (Figure 1 - figure supplement 2A-2D). Of note, the Cre recombinase could be randomly activated in neural epithelial (NE) cells of *Kdm2b^CreERT2/+^;Ai14* mice in the absence of tamoxifen, thus confounding the analysis of lineage-tracing data (Figure 1 - figure supplement 2E-2F). Nonetheless, by P7, tdTomato-positive cells largely express SATB2 (63.6 ± 4.8%) and/or CTIP2 (44.3 ± 5.8%) (Figure 1 - figure supplement 2G-2H). To overcome the issue, we electroporated the LoxP-STOP-LoxP-DsRed (pCALNL) reporter plasmid into the E12.5 *Kdm2b^CreERT2/+^* cortices followed by tamoxifen injection six hours after electroporation. In line with genetic lineage-tracing data, the majority DsRed-positive cells express either SATB2 (71.9% ± 1.5%) or CTIP2 (18.2% ± 7.1%) (Figure 1E). The above expression and lineage-tracing results suggest *Kdm2b* and *LncKdm2b* are transiently expressed in differentiating IPCs and freshly born PNs and might regulate neuronal differentiation during cortical neurogenesis.

### *LncKdm2b* Regulates *Kdm2b’s* Expression *in cis*

The close proximity of *Kdm2b* and *LncKdm2b*’s TSS and their identical expression patterns in developing cortices prompted us to examine if *LncKdm2b* regulates *Kdm2b’s* expression. Since it’s impractical to maintain intermediate progenitor cells or immature projection neurons in *vitro*, we utilized a few primary or immortalized cells that express both *Kdm2b* and *LncKdm2b* to address the issue. First, we transduced Neuro-2a neuroblastoma cells with *LncKdm2b* antisense oligonucleotides (ASOs), which mediate RNA degradation *via* the RNase H-dependent mechanism (Vickers et al., 2003; Walder and Walder, 1988). The levels of *Kdm2b’s* transcripts and protein were significantly decreased upon the ASO treatment (Figure 2A-2B). Consistently, knockdown of *LncKdm2b* by ASO or shRNAs in adherent cultured cortical cells leads to decreased *Kdm2b* expression (Figure 2C, Figure 2 - figure supplement 1A-1B). Next, we applied the CRISPR/Cas9 technique to delete the genomic region of *LncKdm2b’s* second exon in NE-4C mouse neural stem cells (*LncKdm2b^exon2-KO^*), which results in compromised expression of *LncKdm2b* and *Kdm2b* (Figure 2D, Figure 2 - figure supplement 1D). Notably, there’re significant amounts of transcripts derived from *LncKdm2b’s* first and third exons in LncKdm2b^exon2-KO^ cells (Figure 2D). However, the expression levels of *Zfp292*, the downstream target of *LncKdm2b* in ILC3 cells (Liu et al., 2017), were not decreased upon *LncKdm2b* depletion, suggesting a cell-type-specific effects by *LncKdm2b* (Figure 2 - figure supplement 1C, Figure 2 - figure supplement 1E). Therefore, *LncKdm2b* maintains *Kdm2b*’s expression in neural cells.

**Figure 2.**
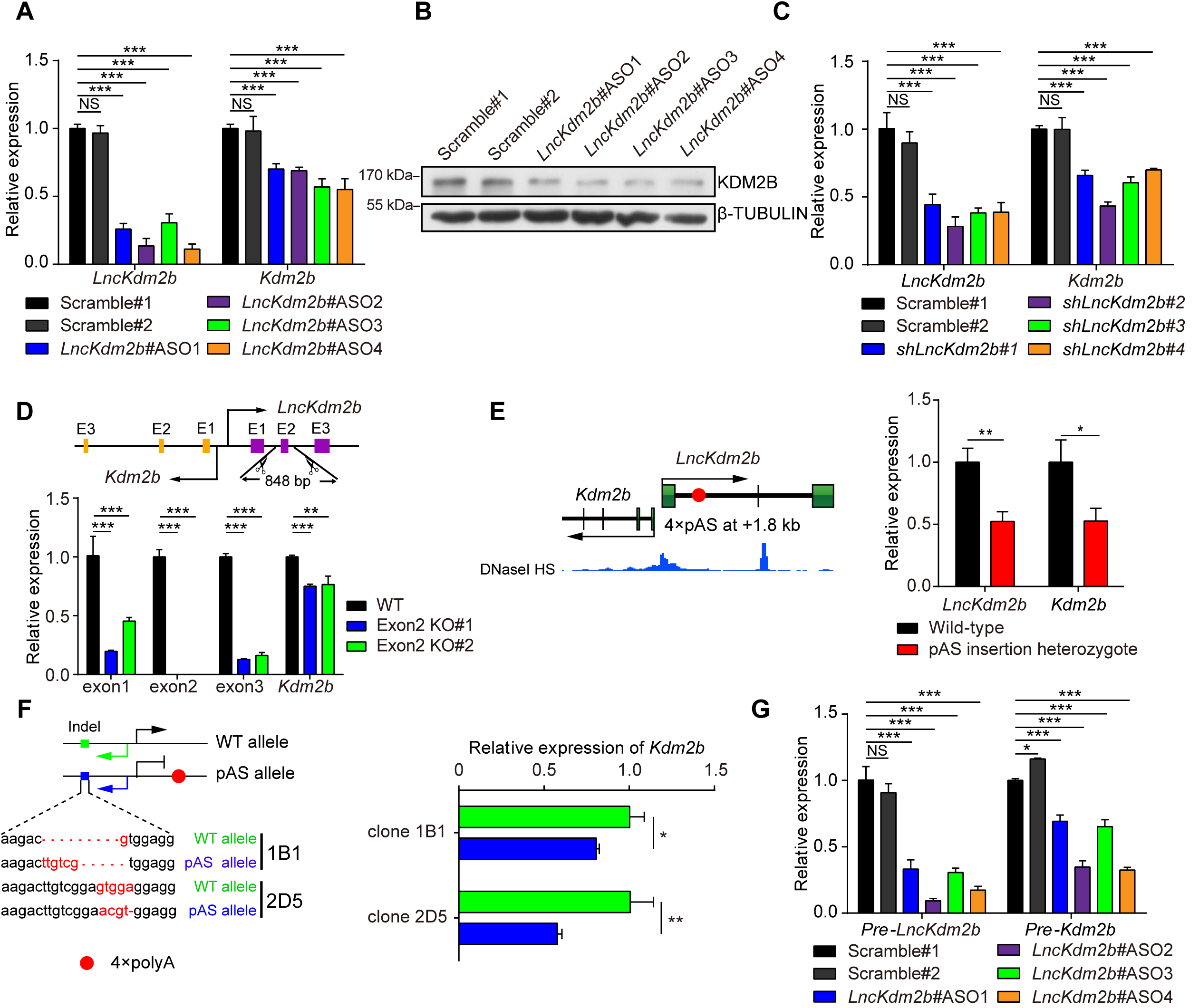
*LncKdm2b* maintains *Kdm2b* Transcription in *cis*. (A) RT-qPCR analysis of *LncKdm2b* and *Kdm2b* RNA levels in Neuro-2a cells treated for two days with Scramble ASOs or ASOs targeting *LncKdm2b*. (B) Representative immunoblotting of Neuro-2a cells treated for four days with indicated ASOs using antibody against KDM2B and β-TUBULIN. (C) RT-qPCR analysis of *LncKdm2b* and *Kdm2b* RNA levels in adherent cultures derived from E12.5 cortices. The cultures were treated with indicated shRNAs. (D) RT-qPCR analysis of *LncKdm2b* and *Kdm2b* RNA levels in wild-type or *LncKdm2b’s* exon 2 knockout NE-4C cells. The expression levels of individual exons of *LncKdm2b* were examined. (E) Left: Schematic diagram showing the insertion of a pAS cassette at 1.8 kb downstream the TSS of *LncKdm2b* in *mESC^LncKdm2b-pASI+^*. pAS, 3 × SV40 polyA and a BGH polyA signal. Right: RT-qPCR analysis of *Kdm2b* mRNA levels in *mESC^LncKdm2b-pAsi^+* and wild-type mESCs. (F) Left: Schematic diagram showing the indels of *Kdm2b’s* second exon in two *mESC^LncKdm2b-pASI+^* clones, 1B1 and 2D5. Right: RT-qPCR analysis of *Kdm2b’s* expression levels from individual alleles of clone 1B1 and 2D5. The y-axis represents relative expression normalized to genomic DNA. (G) The effects of *LncKdm2b* knockdown on nascent transcripts in nuclear run-on assays. RT-qPCR analysis of *LncKdm2b*, and *Kdm2b* nascent transcripts in Neuro-2a cells treated for two days with scramble ASO or ASOs targeting *LncKdm2b*. The y-axis represents relative expression normalized to *Gapdh* nascent transcript. In (A), (C), and (D-G), quantification data are shown as mean + SD (n = 3 unless otherwise indicated). In (A), (C-D), and (G), statistical significance was determined using 2-way ANOVA followed by the Bonferroni’s post hoc test. In (E-F), statistical significance was determined using unpaired 2-tailed Student’s t test. * p<0.05, ** p<0.01, *** p<0.001, “NS” indicates no significance. The y-axis represents relative expression normalized to *Gapdh* transcript unless otherwise indicated. See also Figure 2 - figure supplement 1.

Cross-talk among neighboring genes could involve *trans-* and/or *cis*-regulatory mechanisms, the latter including enhancer-like activity of gene promoters, the process of transcription, and the splicing of the transcript (Bassett et al., 2014; Engreitz et al., 2016; Yin et al., 2015). To discriminate these possibilities, four polyadenylation sequences (pAS) were inserted 1.8 kb downstream of *LncKdm2b’s* TSS to prematurely terminate its transcription in one allele of mouse C57/B6 embryonic stem cells (*mESC*^*LncKdm2b-pAS/+*^), but to keep undisturbed the essential promoter region for *Kdm2b* and *LncKdm2b*’s transcription, which is DNase I hypersensitive (HS) (Figure 2E). Consistently, the expressions of *LncKdm2b* and *Kdm2b* were significantly decreased upon pAS insertion (Figure 2E), suggesting *LncKdm2b’s* transcription process and/or transcripts themselves are required for *Kdm2b’s* expression. We next studied if *LncKdm2b* maintains *Kdm2b’s* transcription *in cis*. First, subcellular fractionation followed by RT-qPCR and RNA ISH assays revealed that most *LncKdm2b* resides in the cytosol with a fraction in the nuclei of cortical cells (Figure 2 - figure supplement 1F-1G). Next, we genetically modified *mESC^LncKdm2b-pAS/^+* cells so that indels were created in the second exon of *Kdm2b* in an allele-specific manner. Quantitative RT-PCR experiments of the two clones (1B1 and 2D5) showed it’s the allele with pAS insertion has significantly lower *Kdm2b* transcription than the other allele (Figure 2F). Lastly, nuclear extracts from *LncKdm2b-depleted* Neuro-2a cells were collected and subjected to nuclear run-on assay. Data showed depletion of *LncKdm2b* results in significantly lower yield of *Kdm2b* nascent transcripts (Figure 2G). In contrast, overexpressing *LncKdm2b in trans* didn’t elevate *Kdm2b’s* transcripts’ levels (Figure 2 - figure supplement 1H). In summary, these results validate *LncKdm2b’s* role in maintaining *Kdm2b* transcription *in cis*.

### *LncKdm2b* Modulates the Configuration of *Kdm2b’s* C/s-regulatory elements

Specific gene expression is coordinated by *cis*-regulatory elements such as the promoter/enhancer, cell-type-specific transcription factors and chromatin states (Heintzman et al., 2009; Perino and Veenstra, 2016). To understand these mechanisms underlying *Kdm2b* transcription, we first analyzed the genomic region both upstream and downstream of the *Kdm2b* and *LncKdm2b’s* TSS. This genome region contains multiple active and/or repressive epigenetic modifications including DNase I HS, H3K27ac (indicative of active enhancers), H3K4me1 (active or poised enhancers), H3K27me3 (repressive or poised *cis*-elements), and CTCF-association (insulators) in developing mouse brain (Vierstra et al., 2014; Yue et al., 2014), suggesting it may contain putative *cis*-regulatory sequences (enhancers) for *Kdm2b* (T1 to T7, Figure 3A). Since **cis*-* regulatory elements/enhancers can be recruited spatially adjacent to promoters to control gene expression, we performed the chromosome conformation capture (3C) followed by qPCR experiments and identified a peak of high crosslinking frequency at the H3K4me1-enriched T5 locus (5.9 kb upstream of *Kdm2b’s* TSS) when using a constant EcoR I fragment located close to the *Kdm2b’s* promoter (Figure 3B), indicating the T5 locus is significantly associated with *Kdm2b’s* promoter. Interestingly, depletion of *LncKdm2b* significantly attenuated the association between T5 and *Kdm2b’s* TSS, suggesting transcribed *LncKdm2b* maintains *Kdm2b’s* expression by inducing a local 3D chromatin structure to bring close *Kdm2b’s* enhancer and promoter (Figure 3B). In addition, luciferase (Luc) assays revealed that the 1.67 kb-long DNA fragment containing the T5 locus has strong enhancer/promoter activities in Neuro-2a cells when it was reversely placed (opposite of *Kdm2b’s* transcription direction) at 5’ of the firefly Luc cassette (Figure 3C). We further narrowed the T5 locus to an evolutionarily conserved 484 bp-long region (T5-mini) and revealed this fragment can also significantly drive Luc expression if reversely placed at 5’ of the firefly Luc cassette. In line with the T5 locus being an evolutionarily conserved *cis*-regulatory element, both mouse T5 and T5-mini sequences are able to drive reporter gene expression in human HEK293T cells (Figure 3 - figure supplement 1A).

To validate if the genomic region embedded with the T5 element is sufficient to initiate spatiotemporal transcription in cortices, we cloned a piece of 8.0 kb genomic DNA (*KUS* – *Kdm2b* upstream sequence, −0.6 kb to +7.3 kb relative to *Kdm2b’s* TSS) from the mouse genome. *In utero* electroporation assay revealed that this genomic region alone can efficiently drive the expression of short-lived d2EGFP (Corish and Tyler-Smith, 1999) in embryonic cortices at either orientation with a pattern reminiscent of endogenous *Kdm2b* or *LncKdm2b* (Figure 3D). In addition, we found the T5 region is essential for *Kdm2b’s* expression, as genomic deletion of T5 leads to compromised *Kdm2b’s* expression in NE-4C cells and in cortical cells (Figure 3E-3F, Figure 3 - figure supplement 1B-1D). Together, these data indicate the *Kdm2b’s* upstream region contains evolutionarily conserved *cis*-regulatory elements essential for expression of *Kdm2b* and *LncKdm2b*, and its configuration is modulated by *LncKdm2b*.

**Figure 3.**
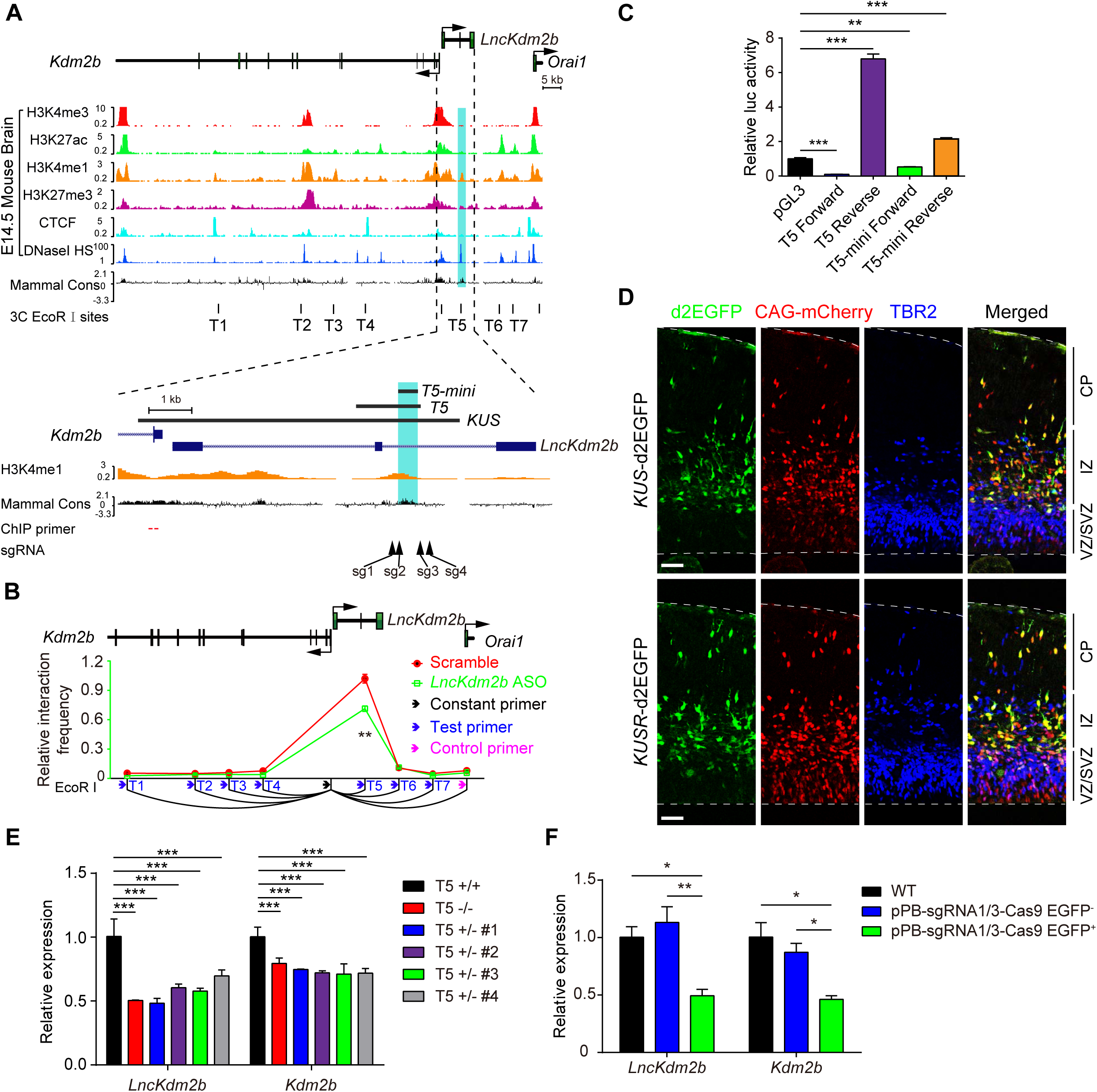
*LncKdm2b* Regulates the Configuration of *Kdm2b’s cis*-elements. (A) Schematic illustration of the *LncKdm2b/Kdm2b* locus. The top tracks show ChIP-seq signals of H3K4me3, H3K27ac, H3K4me1, H3K27me3 and CTCF; and DNase I hypersensitivity (HS) in E14.5 mouse brain, along with sequence conservation among mammals. Bottom tracks show a higher-magnification view of the genomic region covering the promoter for *Kdm2b* and its upstream region that transcribes *LncKdm2b*. Indigo box indicates the conservative region T5 that is also enriched with H3K4me1. KUS, *Kdm2b* upstream sequence. T1 to T7 marks putative regulatory *cis*-elements. (B) Relative crosslinking frequency measured in Neuro-2a cells by 3C-qPCR using a constant primer in an EcoR I fragment at the *Kdm2b* TSS. Cells were treated with Scramble ASO or a mix of ASOs targeting *LncKdm2b* (ASO 1, 2, 3, 4) for two days. Crosslinking frequency is relative to a negative region (the magenta arrow). (C) Luciferase activities in experiments where indicated vectors were transfected into Neuro-2a cells for 24 hours. ‘Forward’ and ‘Reverse’ indicate directions same as or opposite to *Kdm2b’s* transcription orientation. (D) E13.5 mouse cortices were electroporated with KUS-d2EGFP or KUSR-d2EGFP, along with CAG-driving mCherry-expressing vectors. Embryos were sacrificed at E15.5, followed by TBR2 immunofluorescent stainings on coronal cortical sections. Scale bars, 50 μm. KUSR: *Kdm2b* upstream sequence, reversed. (E) RT-qPCR analysis of *LncKdm2b* and *Kdm2b* RNA levels in NE-4C cells with the T5 region knocked out. (F) RT-qPCR analysis of *LncKdm2b* and *Kdm2b* RNA levels in cortical cells with the T5 region knocked out. EGFP+ cells express gRNAs and the Cas9 protein. In (B), quantification data are shown as mean ± SD (n = 3). In (C), (E), and (F), quantification data are shown as mean + SD (n = 3). In (B), statistical significance was determined using 2-tailed Student’s t test. In (C), statistical significance was determined using 1-way ANOVA with Tukey’s post hoc tests. In (E) and (F), statistical significance was determined using 2-way ANOVA followed by the Bonferroni’s post hoc test. * *p*<0.05, ** *p*<0.01, *** *p*<0.001, “NS” indicates no significance. The y-axis represents relative expression normalized to *Gapdh*. See also Figure 3 - figure supplement 1.

### *LncKdm2b* Facilitates a Permissive Chromatin Environment for *Kdm2b’s* Expression by Associating with hnRNPAB

In order to test whether *LncKdm2b* displays intrinsic ability to promote gene expression, we used the Gal4-λN/BoxB system to tether this lncRNA to a heterologous reporter promoter (Figure 4 - figure supplement 1A) (Li et al., 2013; Trimarchi et al., 2014; Wang et al., 2011a). The data showed the full-length *LncKdm2b* and its evolutionarily conserved 5’ part (1-908 nt, transcribed from *LncKdm2b* gene’s first and second exons) could enhance luciferase activities in a dosage-dependent manner, whereas its less conserved 3’ part (909-1896 nt) couldn’t (Figure 4 - figure supplement 1B-1D). Therefore, the 5’ conserved part of *LncKdm2b’s* transcript bears intrinsic-activating function. *LncKdm2b’s* intrinsic-activating capability could be due to its association with *trans*-factor(s). We carried out RNA pull-down experiments using biotinylated *LncKdm2b* and *antisense-LncKdm2b* RNAs. RNA pull-down assay was performed using nuclear protein extracts from cortical NPCs followed by mass spectrometry (MS). A number of RNA binding proteins were enriched in *LncKdm2b*-precipitating extracts compared to those precipitated by antisense *LncKdm2b* (Supplementary file 1 - Table 2). One of the most enriched protein is heterogeneous nuclear ribonucleoprotein A/B (hnRNPAB), which is validated by RNA pull-down followed by immunoblotting (Figure 4A-4B). Notably, SATB1, the protein partner of *LncKdm2b* in group 3 innate lymphoid cells (ILC3) cells (Liu et al., 2017), was not identified to be associated with *LncKdm2b* in this study, probably due to cellular specificity. HnRNPAB is dynamically expressed during brain development and has implications in neuronal differentiation (Sinnamon et al., 2012). Depletion of *Hnrnpab* in Neuro-2a cells significantly decreased *Kdm2b’s* expression (Figure 4 - figure supplement 1E). In control experiments, knockdown the expression of *Dhx9, Satb1, Bptf, Hnrnpa2b1, Hnrnpa3, Dhx5, Ncl* or *Lmnb1*, genes encoding other putative *LncKdm2b*-associated proteins, had no significant effect on *Kdm2b’s* expression (Figure 4 - figure supplement 1E). RNA *in situ* hybridization followed by immunofluorescent staining showed colocalization of *LncKdm2b* and hnRNPAB in cortical NPCs (Figure 4 - figure supplement 1F). RNA immunoprecipitation experiments (RIP) in either native or the formaldehyde-fixed condition confirmed association of hnRNPAB with *LncKdm2b* but not with *Actb* or *Gapdh* RNAs (Figure 4C-4D).

**Figure 4.**
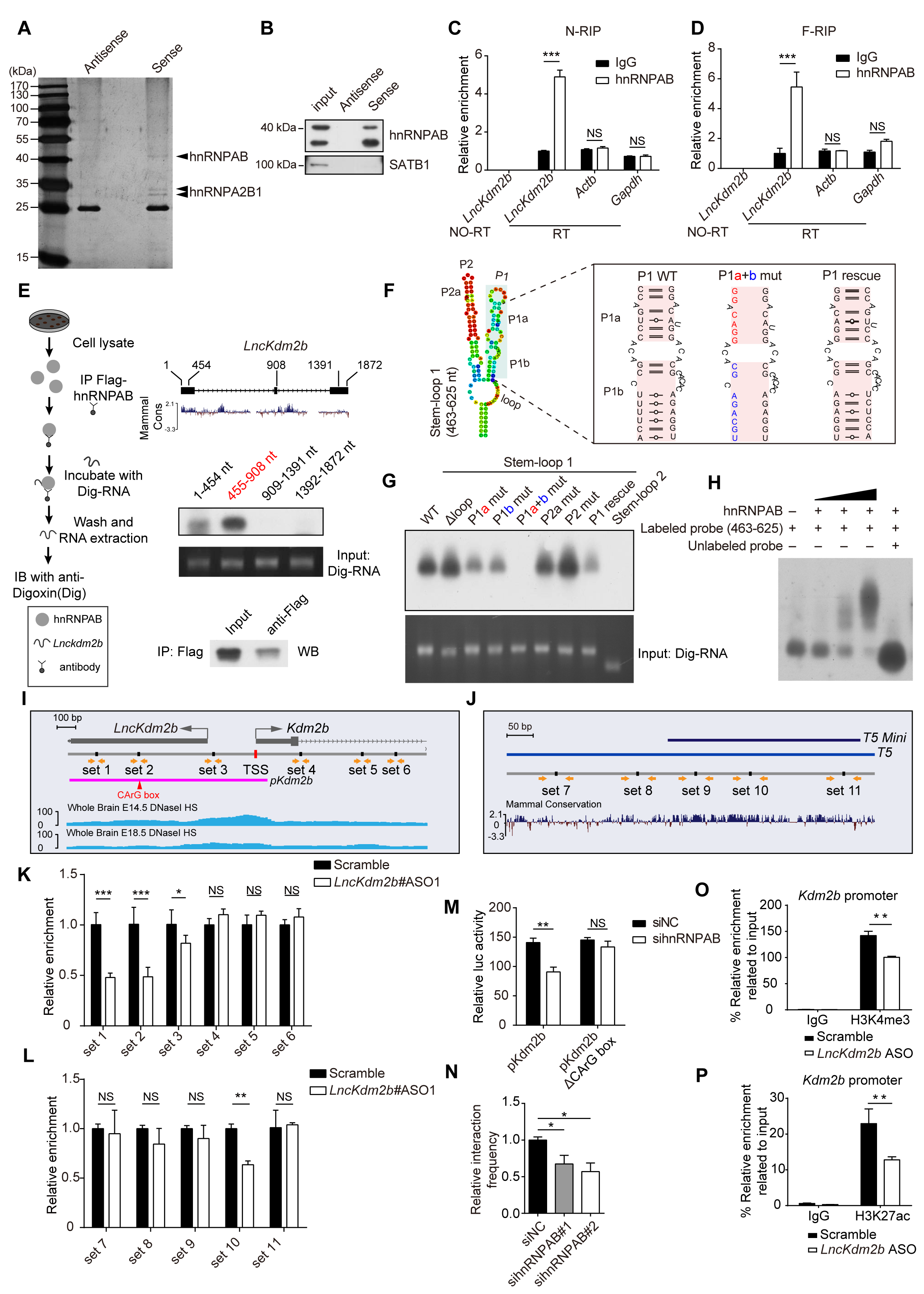
*LncKdm2b* Transcripts Can Activate Gene Expression and Modulate Local Chromatin State. (A) Identification of proteins associated with *LncKdm2b*. Protein extracts from E14.5 mouse cortices were incubated with the biotinylated *LncKdm2b* (sense) or control (antisense *LncKdm2b*) followed by SDS-PAGE and silver staining. (B) Immunoblots of hnRNPAB and SATB1 of protein extracts that are associated with sense or antisense *LncKdm2b* in Neuro-2a cells. (C-D) RNA immunoprecipitation (RIP) of anti-hnRNPAB and control IgG antibodies in native (N-RIP, C) and formaldehyde treated (F-RIP, D) Neuro-2a cells. Extracted RNAs were subjected to RT-qPCR analysis of indicated transcripts. (E) Digoxigenin-labeled *LncKdm2b* truncations were incubated with Flag-hnRNPAB bound to anti-Flag-agarose beads. hnRNPAB associated RNAs were chemiluminescently detected. (F) Left: schematic diagram showing the putative stem-loop structure in *LncKdm2b’s* 455908 nt region. Right: point mutations made to disrupt hairpin formation. (G) Digoxigenin-labeled *LncKdm2b’s* stem loops described in (F) were incubated with Flag-hnRNPAB bound to anti-Flag-agarose beads. hnRNPAB associated RNAs were chemiluminescently detected. (H) EMSA assays of digoxigenin-labeled *LncKdm2b* RNA (463-625 nt) incubated with purified hnRNPAB. (I-J) Schematic illustration of primer sets used in ChIP-qPCR experiments around the TSS of the *LncKdm2b/Kdm2b* locus (I) and the T5 region (J). *Kdm2b’s* promoter region (*pKdm2b*) and the putative hnRNPAB-binding CArG box for luciferase reporter assay in (M) was also shown. (K-L) ChIP-qPCR analysis of indicated primer sets showed in (I-J) enriched by anti-hnRNPAB antibody upon depletion of *LncKdm2b*. The y-axis shows fold enrichment normalized to scramble ASO control. (M) Relative luciferase activity of T5 region with or without the CArG box in Neuro-2a cells. Cells were treated for two days with siRNAs against hnRNPAB. (N) Relative crosslinking frequency between the T5 and *Kdm2b’s* TSS upon hnRNPAB depletion measured by 3C-qPCR in Neuro-2a cells. The y-axis shows fold enrichment normalized to the scramble control (siNC). (O-P) Neuro-2a cells were treated with Scramble ASO or a mix of ASOs targeting *LncKdm2b* (ASO 1, 2, 3, 4) for 48 hours before ChIP-qPCR of H3K4me3 (O), and H3K27ac (P) at the *Kdm2b* promoter. The y-axis shows fold enrichment normalized to the input. Positions of promoter primers are shown on the bottom of Figure 3A. Quantification data are shown as mean + SD (n = 3). In (C-D), (K-M), and (O-P), statistical significance was determined using 2-tailed Student’s t test. In (N), statistical significance was determined using 1-way ANOVA with Tukey’s post hoc tests. * *p*<0.05, ** *p*<0.01, *** *p*<0.001, “NS” indicates no significance. See also Figure 4 - figure supplement 1.

In line with the fact that the 5’ conserved part of *LncKdm2b* has intrinsic-activating function (Figure 4 - figure supplement 1B-1D), *in vitro* binding experiments indicated the two 5’ conserved regions (1-454 nt and 455-905 nt) of *LncKdm2b* could interact with hnRNPAB, with the 455-905 nt region having stronger association with hnRNPAB than the 1-454 nt region. On the other hand, the 3’ non-conserved region (909-1391nt and 1392-1872 nt) couldn’t associate with hnRNPAB (Figure 4E). RNA structure analysis using *RNAfold* predicts two stem-loops in the 455-905 nt region (Figure 4 - figure supplement 1G). Particularly, the stem-loop 1 (463-625 nt) has two hairpin arms, P1 and P2 (Figure 4F). To ask if these hairpin arms are required for the interaction between *LncKdm2b* and hnRNPAB, we mutated a few nucleotides to disrupt the hairpin formation (Figure 4F). In *vitro* binding experiments indeed showed disruption of the hairpin formation in P1 would greatly compromised the interaction (Figure 4G). Moreover, restoration of the P1 hairpin (P1 rescue) would partially rescue the association, but the P2 hairpin or the stem-loop 2 (840-918 nt) is not required for the association of *LncKdm2b* with hnRNPAB. The EMSA (electrophoretic mobility shift assay) experiment further validated the binding of *LncKdm2b’s* stem-loop 1 region (463-625nt) to hnRNPAB (Figure 4H). Together, these analyses revealed that the hairpin P1 of *LncKdm2b*’s conserved 5’ part directly interacts with hnRNPAB, which might be responsible for *LncKdm2b’s* intrinsic-activating function (Figure 4 - figure supplement 1B-1D).

HnRNPAB, also known as CArG box-binding factor-A (CBF-A), is an RNA binding protein with transcription activity (Venkov et al., 2007; Zhou et al., 2014). We went on to ask if hnRNPAB binds to genomic regions essential for *Kdm2b* expression, and if the binding is regulated by *LncKdm2b*. ChIP-qPCR showed hnRNPAB binds to multiples sites in the *Kdm2b’s* promoter and the T5 region, many of which were positively regulated by *LncKdm2b* (Figure 4 - figure supplement 1H, Figure 4I-4L). The reporter activity driven by *Kdm2b’s* promoter (*pKdm2b)* is mediated by the hnRNPAB-binding CArG box. Downregulating hnRNPAB would significantly lower *pKdm2bs* reporter activity, whereas exert no effect on CArG box-deleted *pKdm2b* (Figure 4M). Moreover, the association between the T5 and *Kdm2b’s* TSS was significantly compromised upon hnRNPAB depletion (Figure 4N). Finally, the *Kdm2b’s* promoter (−78 bp to −20 bp relative to *Kdm2b’s* TSS) was less enriched for H3K4me3 and H3K27ac, two histone markers indicative of active transcription, in *LncKdm2b-depleted* Neuro-2a cells (Figure 4O-4P). Collectively, *Kdm2b’s* expression correlates positively with the association between *Kdm2b’s* promoter and an essential enhancer (T5), which is facilitated by *LncKdm2b’s* transcripts and its associated protein hnRNPAB. These findings point a role of *LncKdm2b* in regulating transcription locally.

### KDM2B Promotes Cortical Neuronal Differentiation

Since *LncKdm2b* regulates the expression of *Kdm2b*, and *Kdm2b* is transiently expressed in freshly born projection neurons, we next explored roles and mechanisms of KDM2B in cortical neurogenesis. We first electroporated E13.5 embryonic cortices with plasmids overexpressing *Kdm2b* and collected brains at E15.5 (Figure 5 - figure supplement 1A). Significantly more *Kdm2b* transduced cells reside in the cortical plate (CP, future cortices) with fewer cells in the VZ/SVZ, indicating accelerated cortical neurogenesis and radially neuronal migration (Figure 5A-5B). In line with this, fewer mCherry+ *Kdm2b* transduced cells express TBR2 and PAX6, markers for IPCs and RGPCs respectively (Figure 5C-5E). Embryonic brains of *Kdm2b^CreERT2/CreERT2^* mice have significant amount of residual KDM2B protein probably due to inefficient transcriptional termination (Figure 5 - figure supplement 1B), which might lead to subsequent use of alternative start codons. We therefore performed *Kdm2b* loss-of-function studies by electroporating plasmids expressing short-hairpin RNAs (shRNAs) against *Kdm2b* into E13.5 embryonic cortices. To minimize nonspecific effects, we chose the shmiRNA system to express long RNA hairpins with shRNAs embedded into endogenous miRNA loop and flanking sequences (Baek et al., 2014; Bauer et al., 2009). Significant more *Kdm2b*-shRNA electroporated cells reside in the VZ/SVZ at E16.5 (Figure 5F-5G). Next, E16.5 *Kdm2b*-shRNA transduced cortices were immuno-stained with TBR2 and NEUROD2, a transcriptional factor expressed in cortical PNs. Results showed more transduced cells are co-labeled with TBR2 but fewer cells express NEUROD2, with significantly more NEUROD2+ transduced cells localized in the VZ/SVZ (Figure 5H-5I, Figure 5 - figure supplement 1C-1D). Moreover, more transduced cells are colocalized with PAX6-positive RGPCs (Figure 5J-5K). This phenotype can be fully rescued by simultaneously overexpressing *Kdm2b* (Figure 5F-5G, Figure 5 - figure supplement 1E-1F). Furthermore, significantly more *Kdm2b*-depleted cells (EGFP+) are BrdU positive and in S-phase, as embryos were injected BrdU 30 minutes before sacrifice; and more PAX6+EGFP+ RGPCs are BrdU positive, suggesting depletion of *Kdm2b* promotes proliferation of RGPCs (Figure 5L-5N). We didn’t observed changes of programmed cell death (cleaved caspase-3+ cells) in *Kdm2b*-shRNA transduced cortices (Figure 5 - figure supplement 1G). All these data support the notion that KDM2B promotes cortical neuronal differentiation in *vivo* (Supplementary file 1 - Table 3).

**Figure 5.**
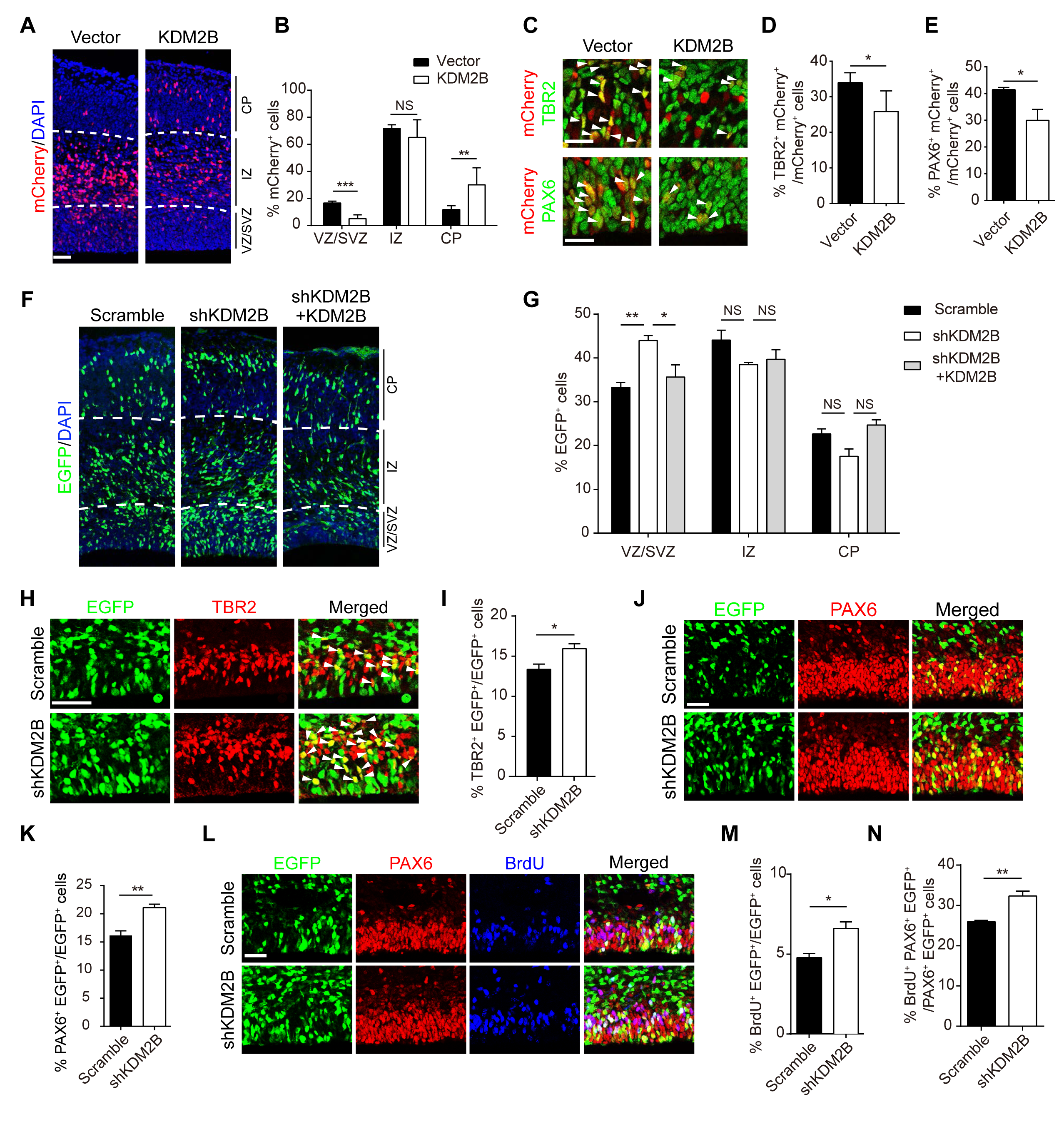
*Kdm2b* promotes Cortical Neurogenesis. (A-E) E13.5 mouse cortices were electroporated with empty or *Kdm2b*-expressing vector, along with mCherry-expressing vector to label transduced cells. Embryos were sacrificed at E15.5 for immunofluorescent analysis. Coronal Sections were stained for DAPI (A), and the relative location of mCherry-positive cells was quantified (B). Ten embryos in control and nine embryos in *Kdm2b*-overexpression. Representative VZ/SVZ images of control and *Kdm2b*-expressing cortices immunostained for TBR2 (top) and PAX6 (bottom). Arrowheads denote double-labeled cells (C). Quantification of TBR2+ (D) or PAX6+ (E) cells in transduced cells. (F-N) E13.5 mouse cortices were electroporated with indicated combination of vectors, with transduced cells labeled with EGFP. Embryos were sacrificed at E16.5 for immunofluorescent analyses. The relative location of EGFP+ cells was quantified (F-G). Three embryos in scramble, and shKDM2B, five embryos in shKDM2B plus KDM2B. Representative VZ/SVZ images of scramble or KDM2B shRNA electroporated sections immunostained for TBR2 (H) and quantification of TBR2+ transduced cells (I). Representative VZ/SVZ images of scramble or KDM2B shRNA electroporated sections immunostained for PAX6 (J) and quantification of PAX6+ transduced cells (K). Sections were co-immunostained with PAX6 and BrdU (30 minutes) (L). Percentiles of BrdU+ transduced cells (M), and of BrdU+PAX6+EGFP+ / PAX6+EGFP+ cells (N). In (B), (D-E), (G), (I), (K), and (M-N), quantification data are shown as mean + SEM. In (B), (D-E), (I), (K), and (M-N), statistical significance was determined using 2-tailed Student’s t test. In (G), statistical significance was determined using 2-way ANOVA followed by the Bonferroni’s post hoc test.* *p*<0.05, ** *p*<0.01, *** *p*<0.001, “NS” indicates no significance. Scale bars, 50 μm. VZ, ventricular zone; SVZ, subventricular zone; IZ, intermediate zone; CP, cortical plate. See also Figure 5 - figure supplement 1.

### KDM2B Depends on its Leucine-rich Repeats to Promote Cortical Neuronal Differentiation

KDM2B contains multiple functional domains, including the JmjC histone demethylase domain, a DNA-binding CxxC zinc finger, an F-box domain, a PHD finger, and eight leucine-rich repeats (LRRs). The F-box and the LRR domain are required for variant polycomb repressor complex 1 (PRC1) recruitment and assembly (Farcas et al., 2012; He et al., 2013; Inagaki et al., 2015; Wu et al., 2013). To decipher how KDM2B exerts its function in cortical development, we carried out rescue experiments using *in utero* electroporation (Figure 6A-6D). E13.5 cortices were co-electroporated with plasmids expressing shRNA against *Kdm2b* along with plasmids expressing different KDM2B truncations or mutations (Figure 6 - figure supplement 1A). Intriguingly, expressing KDM2B with JmjC mutation (lacking methyl-transferase activity), PHD mutation; CxxC, or F-box deletion individually could fully rescue hampered radial migration (judged by relative positions of transduced cells) and enhanced RGPCs self-renewal (determined by PAX6+ transduced cells) caused by *Kdm2b* depletion, whereas the LRR-deleted KDM2B (KDM2B-ΔLRR) could not rescue the defect (Figure 6A-6D). Moreover, overexpressing the *Kdm2b*-ΔLRR alone also leads to delayed neuronal differentiation and radial migration, and enhanced self-renewal of RGPCs (Figure 6E-6G), suggesting the LRR domain is indispensable for KDM2B’s role in promoting cortical neuronal differentiation and *Kdm2b*-ΔLRR overexpression caused a dominant-negative effect (Supplementary file 1 - Table 3).

**Figure 6.**
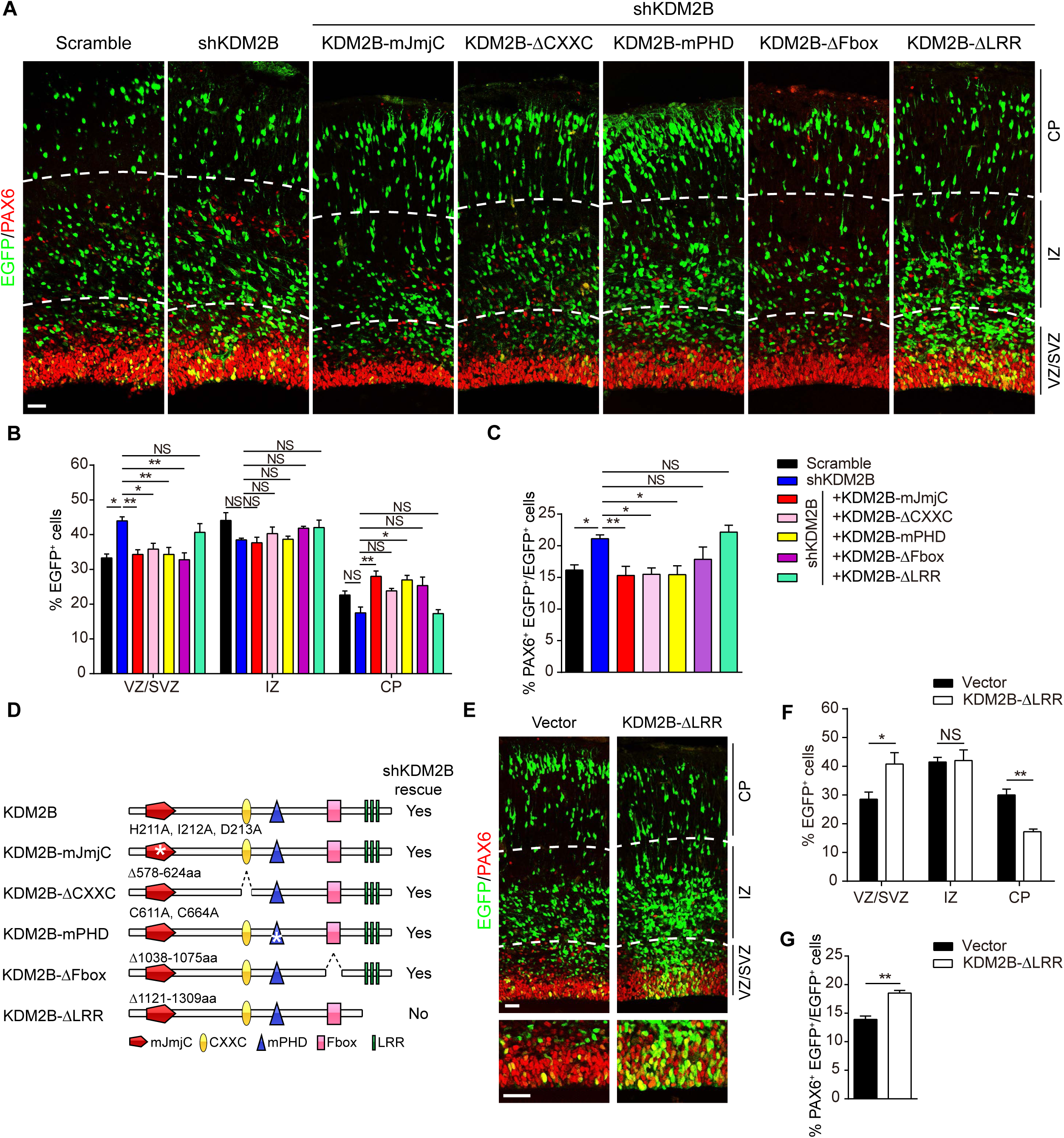
KDM2B relies on its leucine-rich repeats (LRR) to promote cortical neurogenesis. (A-D) E13.5 mouse cortices were electroporated with indicated combinations of vectors, with transduced cells labeled with EGFP. Embryos were sacrificed at E16.5 for PAX6 immunofluorescent stainings (A). Three embryos in scramble and KDM2B shRNA, six embryos in KDM2B shRNA plus *Kdm2b*-mJmjC and KDM2B shRNA plus *Kdm2b*-ΔCxxC, five embryos in KDM2B shRNA plus *Kdm2b*-mPHD, four embryos in KDM2B shRNA plus *Kdm2b*-ΔFbox, and five embryos in KDM2B shRNA plus *Kdm2b*-ΔLRR. The relative location of EGFP+ cells (B) and percentiles of PAX6+ transduced cells (C) were quantified. Illustration showing KDM2B constructs for rescue assays (D). (E-G) E13.5 mouse cortices were electroporated with empty or *Kdm2b*-ΔLRR-expressing vector, along with EGFP-expressing vectors to label transduced cells. Embryos were sacrificed at E16.5 for PAX6 immunofluorescent analysis (E). The relative location of EGFP+ cells (F) and percentiles of PAX6+ transduced cells (G) were quantified. Six embryos each. In (B-C) and (F-G), quantification data are shown as mean + SEM. In (B), statistical significance was determined using 2-way ANOVA followed by the Bonferroni’s post hoc test. In (C), statistical significance was determined using 1-way ANOVA with Tukey’s post hoc tests. In (F-G), statistical significance was determined using 2-tailed Student’s t test. * *p*<0.05, ** *p*<0.01, *** *p*<0.001, “NS” indicates no significance. Scale bars, 50 μm. VZ, ventricular zone; SVZ, subventricular zone; IZ, intermediate zone; CP, cortical plate. See also Figure 6 - figure supplement 1.

It has been reported that the LRR domain is essential for KDM2B to recruit and assemble the PRC1; and depletion of RING1B, the catalytic component of PRC1, in mice leads to prolonged neurogenic phase of NPCs and delaying of the onset of the astrogenic phase (Hirabayashi et al., 2009; Morimoto-Suzki et al., 2014). To study if KDM2B relies on PRC1 to exert its role, we electroporated plasmids expressing catalytic inactive RING1B (I53A) into the E13.5 cortices. Although this mutation ablates RING1B’s ability to act as an E3 ligase *in vitro* (Buchwald et al., 2006), it does not perturb the incorporation of RING1B into canonical and variant PRC1 (Illingworth et al., 2012). Surprisingly, the neurogenic process is unaltered by E16.5 (Figure 6 - figure supplement 1B-1E). Moreover, whole-body inactivation of *Kdm2b* in mice doesn’t lead to reduction of H2AK119Ub1, a histone modification mediated by PRC1 complex (Figure 6 - figure supplement 1F-1G). These results indicate the pro-neurogenic roles exerted by KDM2B is independent of its function in mediating PRC1 activities.

### *LncKdm2b* Promotes Cortical Neuronal Differentiation *via* KDM2B

As we have shown that *LncKdm2b* is transiently expressed in freshly born projection neurons and *LncKdm2b* c/s-activates *Kdm2b* expression, we expected that *LncKdm2b* and *Kdm2b* may have similar function on cortical neuronal differentiation. To this end, we first knocked down the expression of *Kdm2b* or *LncKdm2b* by transfecting adherent-cultured cortical progenitor cells (NPCs) with low titer lentiviral shRNAs to study cell fate changes at the clonal level. NPCs depleted with *Kdm2b* or *LncKdm2b* showed enhanced self-renewal but decreases neuronal differentiation: significantly more *Kdm2b* or *LncKdm2b-depleted* cortical cells expressing SOX2 with fewer cells expressing TUJ1 compared to scramble shRNA-transfected cells (Figure 7A-7B); more precursor-containing clones with fewer neuron-containing clones and fewer TUJ1-only neuronal clones (Figure 7C-7E); and more SOX2+ cells per clone upon *Kdm2b* or *LncKdm2b* depletion (Figure 7F-7G). Thus, *LncKdm2b* and *Kdm2b* are required for proper neuronal differentiation of cortical NPCs in *vitro* (Supplementary file 1 - Table 3).

**Figure 7.**
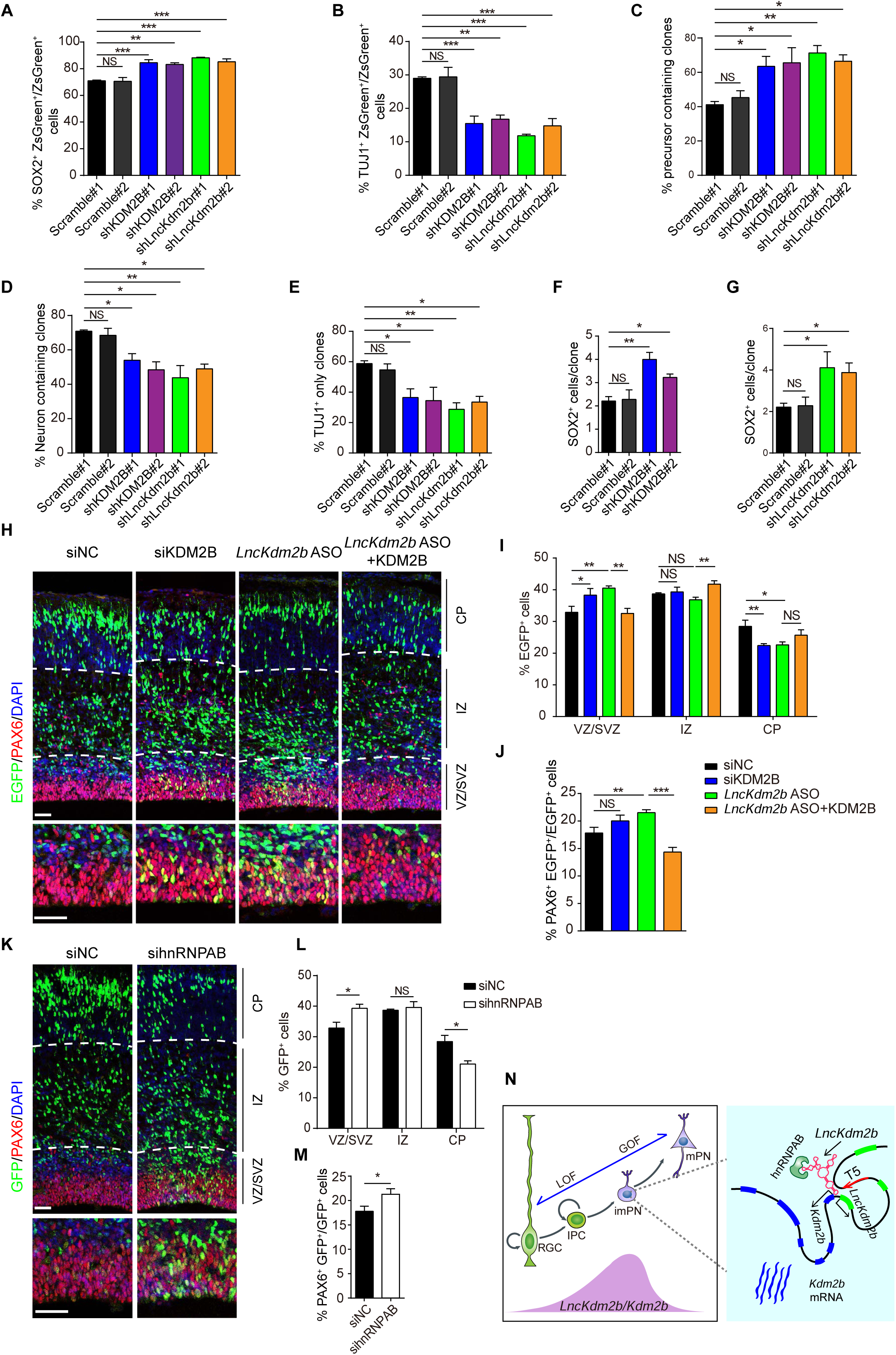
*LncKdm2b* Maintains Mouse Cortical Neurogenesis through KDM2B. (A–G) E12.5 cortical neural precursors were infected with lentivirus expressing indicated shRNAs for three days followed by immunostaining of SOX2 and TUJ1. Transfected cells were labeled with ZsGreen. Quantification analyses were performed to calculate percentiles of SOX2+ (A) or TUJ1+ (B) ZsGreen+ transduced cells; percentiles of clones with at least one SOX2+ precursor (C), clones with at least one TUJ1+ neuron (D), neuron only clones (E); and the average number of SOX2+ cells in SOX2+ clones (F-G). (H-J) E13.5 mouse cortices were electroporated with indicated siRNAs and vectors, with transduced cells labeled with EGFP. Embryos were sacrificed at E16.5 for PAX6 immunofluorescent staining (H). The relative location of EGFP+ cells (I) and percentiles of PAX6+ transduced cells (J) were quantified. Three embryos in control (siNC), five embryos in siKDM2B, *LncKdm2b* ASO, and *LncKdm2b* ASO plus KDM2B. (K-M) E13.5 mouse cortices were electroporated with indicated siRNAs, with transduced cells labeled with EGFP. Embryos were sacrificed at E16.5 for PAX6 immunofluorescent stainings (K). The relative location of EGFP+ cells (L) and percentiles of PAX6+ transduced cells (M) were quantified. Three embryos each. (N) A model for *LncKdm2b* promoting cortical neurogenesis by *cis*-activating *Kdm2b. LncKdm2b* and *Kdm2b* are transiently expressed in freshly born projection neurons. *LncKdm2b* RNA facilitates an open chromatin configuration locally by bringing together the upstream regulatory *cis*-element T5, *Kdm2b* promoter and hnRNPAB to maintain *Kdm2b*’s transcription. In (A-G), (I-J), and (L-M), quantification data are shown as mean + SEM. In (A-G), (J), statistical significance was determined using 1-way ANOVA with Tukey’s post hoc tests. In (I), statistical significance was determined using 2-way ANOVA followed by the Bonferroni’s post hoc test. In (L-M), statistical significance was determined using 2-tailed Student’s t test. * *p*<0.05, ** *p*<0.01, *** *p*<0.001, “NS” indicates no significance. Scale bars, 50 jm. VZ, ventricular zone; SVZ, subventricular zone; IZ, intermediate zone; CP, cortical plate. RGC, radial glial cells; IPC, intermediate progenitor cells; imPN, immature projection neurons; mPN, mature projection neurons; LOF, loss-of-function; GOF, gain-of-function. See also Figure 7 - figure supplement 1.

Next, we explored whether *LncKdm2b* regulates cortical neurogenesis *in vivo* through *Kdm2b*. E13.5 embryonic cortices were electroporated with siRNAs or antisense oligonucleotides (ASO) targeting *Kdm2b* or *LncKdm2b* respectively followed by phenotypic analyses at E16.5. Significantly fewer siKdm2b‐ or *LncKdm2b* ASO-transduced cells reside in the CP with more cells in the VZ/SVZ, indicating delayed neuronal differentiation. In line with this, more transduced cells express PAX6. Most importantly, overexpressing *Kdm2b* can mostly rescue the phenotypes caused by *LncKdm2b* knockdown (Figure 7H-7J). Finally, we ask if hnRNPAB, the *LncKdm2b-* associated protein, also regulates neuronal differentiation in developing neocortex. To this end, we electroporated E13.5 cortices with siRNAs against *Hnrnpab* and indeed found *Hnrnpab-depleted* cells showed delayed neuronal migration to the CP at E16.5 and hampered differentiation of NSPCs - more sihnRNPAB-transduced cells localized in the VZ/SVZ and co-localized with PAX6 (Figure 7K-7M). On the other hand, depletion of *Hnrnpa2b1* didn’t cause such defects (Figure 7 - figure supplement 1A-1C). Moreover, overexpression of *LncKdm2b* has no effect on neuronal migration and differentiation (Figure 7 - figure supplement 1D-1E), which is in line with aforementioned data showing *LncKdm2b* couldn’t trans-activate *Kdm2b* expression (Figure 2 - figure supplement 1H). Together, *LncKdm2b* promotes cortical neuronal differentiation via KDM2B (Supplementary file 1 - Table 3).

In summary, we found the precise balance of self-renewal and neuronal differentiation of NSPCs during cortical neurogenesis is modulated by KDM2B in the LRR-dependent manner. Moreover, the expression of *Kdm2b* is positively regulated by its divergent lncRNA *LncKdm2b*, which facilitates a permissive chromatin configuration locally by bringing together the upstream regulatory *cis*-element T5, *Kdm2b’s* promoter and hnRNPAB (Figure 7N).

## Discussion

The generation of layer-specific PNs over developmental time is precisely controlled and largely attributed to cell-intrinsic properties of NSPCs (Gaspard et al., 2008; Shen et al., 2006). Cell fates choices are mostly the results of specific transcriptional events, which are coordinated by *cis*-regulatory elements, cell-specific transcription factors, and epigenetic states including DNA methylation, histone modification and chromatin accessibility (Heintzman et al., 2009; Perino and Veenstra, 2016). Some lncRNAs can regulate gene transcription locally (*cis)* and/or distally (*trans)* by modifying epigenetic states (Berghoff et al., 2013; Fu, 2014; Grote et al., 2013; Rinn and Chang, 2012). Here we found lncRNA gene *LncKdm2b* shares the same promoter with its bidirectional protein-coding gene *Kdm2b*, and both of them are transiently expressed in committed IPCs and freshly-born PNs. Unlike most bidirectional coding-noncoding transcripts, *LncKdm2b’s* expression level is comparable with that of *Kdm2b* at the peak of cortical neurogenesis, strongly indicating *LncKdm2b’s* regulatory roles. Indeed, *LncKdm2b* maintains *Kdm2b’s* expression *in cis* in neural cells. Mechanistically, the *LncKdm2b* transcripts enhances physical association of *Kdm2b’s* promoter and a key enhancer T5 *via* binding to hnRNPAB. *LncKdm2b’s* transcript, especially its evolutionarily conserved 5’ part, bears intrinsic-activating function and interacts with hnRNPAB *via* one of its putative stem-loop structures (Figure 4A-4H, S4A-S4D). Similarly, a 5’ fragment of *LncKdm2b* (450–700 nt) is necessary for its binding to SATB1 or SRCAP in ILC3 and ES cells respectively (Liu et al., 2017; Ye et al., 2018). HnRNPAB, an RNA binding protein with transcription activity (Venkov et al., 2007; Zhou et al., 2014), was shown to be associated with *Kdm2bs* TSS and the T5 region in neural cells, and the strength of the association depends on the presence of *LncKdm2b* (Figure 4 - figure supplement 1H, Figure 4K-4L). Moreover, the *cis*-activity of *Kdm2b’s* promoter also relies on hnRNPAB’s binding (Figure 4M). The core T5-region (T5-mini), a conserved *cis*-regulatory element embedded in *LncKdm2b’s* second intron, can drive gene expression in both mouse and human cells when reversely placed upstream of reporters, and its deletion results in decreased expression of *Kdm2b*. In summary, this study indicates a role of LncRNA in coordinating the association of *cis*-regulatory elements (*Kdm2b’s* TSS and T5) and *trans*-factor(s) (hnRNPAB) in transcriptional regulation, which probably relies on RNA’s specific secondary structures.

A recent study by Liu *et al*. showed *LncKdm2b* activates expression of *Zfp292 in trans via* recruiting the chromatin organizer SATB1 and the nuclear remodeling factor (NURF) complex onto the *Zfp292* promoter in innate lymphoid cells (ILCs) (Liu et al., 2017). Similarly, *LncKdm2b* activates the expression of *Zbtb3* by promoting the assembly and ATPase activity of the SRCAP complex in mESCs (Ye et al., 2018). Surprisingly, these studies didn’t detect expression alterations of *Kdm2b* in *LncKdm2b-null* ILC3s and mESCs. In our study, *LncKdm2b* was not found to be associated with SATB1. Furthermore, the expression levels of *Zfp292* were not decreased in neural cells depleted with *LncKdm2b* (Figure 2 - figure supplement 1C-1E). These discrepancies could be due to different cellular context and/or distinct inactivation approaches. *LncKdm2b’s* second exon was deleted in previous studies to abolish its transcripts, which might not hamper *LncKdm2b’s* transcription process *per se* and/or *LncKdm2b’s* conserved region with intrinsic-activating function could still exist. In fact, a good fraction of *LncKdm2b’s* transcripts derived from the first and third exons could be detected in NE-4C cells with *LncKdm2b’s* second exon deleted (Figure 2D). In contrast, our study also used siRNAs and ASOs to target *LncKdm2b*, and inserted pAS sites into *LncKdm2b’s* first intron in mESCs, thus impeding *LncKdm2b* transcription, ultimately leading to attenuation of *Kdm2b* transcription (Figure 2A-2G). Interestingly, although *LncKdm2b* controls *Kdm2b’s* expression at the transcriptional level in cell nuclei, a good fraction of *LncKdm2b* transcripts resides in the cytoplasm. Previous studies also indicated *LncKdm2b* localizes in both nuclei and cytosol in mESCs and innate lymphoid cells (Liu et al., 2017; Ye et al., 2018). It remains to be elucidated if *LncKdm2b* functions in cytosol, and if *LncKdm2b*’s cytosolic translocation would facilitate its decay to ensure *Kdm2b*’s transient expression during neuronal differentiation. This finding is just the beginning to understand how LncRNAs regulate cortical neuronal differentiation by controlling local transcription and might have general implications in cell fate determinations.

KDM2B, also known as JHDM1B, NDY1 and FBXL10, was initially characterized as a Jumonji (JmjC) domain containing histone H3K36 di-demethylase. KDM2B is a multidomain protein that is localized to essentially all CpG-rich promoters *via* its CxxC domain that binds to unmethylated CpG dinucleotides. KDM2B is also a constituent of a non-canonical (variant) polycomb repressor complex 1 (PRC1) (Farcas et al., 2012; He et al., 2013; Wu et al., 2013), and KDM2B’s leucine-rich repeat (LRR) domain and the F-box domain are essential for KDM2B to recruit/assemble the PRC1 (Boulard et al., 2015; Inagaki et al., 2015). KDM2B plays pivotal roles in cell senescence and proliferation, DNA repair, embryogenesis, oncogenesis, and somatic cell reprogramming (Andricovich et al., 2016; He et al., 2008; Jiang et al., 2015; Kottakis et al., 2014; Li et al., 2017a; Liang et al., 2012; Wang et al., 2011b). We found KDM2B relies on its LRR domain but not the F-box to regulate neuronal differentiation (Figure 6). Leucine-rich repeats are frequently involved in mediating protein–protein interactions, but most of human LRR-containing proteins remain functionally uncharacterized (Ng and Xavier, 2011). In addition, the E3 ligase activity of RING1B, PRC1’s core component, seems dispensable for cortical neurogenesis (Figure 6 - figure supplement 1C-1E), which is in line with findings showing the enzymatic capabilities of RING1B is dispensable for early mouse development (Illingworth et al., 2015). Consistently, inactivation of *Kdm2b* in mice doesn’t result in reduction of H2AK119Ub1, a histone modification mediated by PRC1 complex (Figure 6 - figure supplement 1F-1G), suggesting KDM2B relies on signals other than PRC1 to promote neuronal differentiation. The deletion or mutation of JmjC, CxxC, PHD domains individually could rescue hampered neuronal differentiation caused by *Kdm2b* loss (Figure 6A-6D), but these domains may still exert scaffold or adaptor roles.

It will be also worthy of exploring how the transient expressions of *Kdm2b* and *LncKdm2b* are initiated and maintained in cortical IPCs and freshly-born PNs. A report showed KDM2B’s expression in primary MEFs and cancer cells is induced by FGF-2 via CREB phosphorylation and activation, downstream of DYRK1A kinase (Kottakis et al., 2011). Since both FGF-2 and DYRK1A have essential roles in cortical development, it remains to be studied if they regulate *KDM2B’s* expression in this context (Arron et al., 2006; Benavides-Piccione et al., 2005; Fotaki et al., 2002; Ghosh and Greenberg, 1995; Vescovi et al., 1993). Nonetheless, we italic *Kdm2b’s* novel function in promoting neuronal differentiation, which is PRC1 independent. Moreover, *Kdm2b’s* expression is maintained by its divergent lncRNA *LncKdm2b*, which mediates a permissive chromatin environment around *Kdm2b’s* promoter. Since normal cortical development is key to neurological functions such as cognition, KDM2B may have implications in neuropsychiatric disorders. In line with this, *KDM2B* is among the most frequently deleted genes in the 12q24.31 microdeletion syndrome, which is characterized by principal clinical features including autism, intellectual disability, epilepsy, and craniofacial anomalies (Labonne et al., 2016). Intriguingly, human *LncKDM2B* is also transcribed divergently from the promoter of *KDM2B* with high sequence homology with *LncKdm2b* (Ye et al., 2018). It remains to be investigated if *LncKDM2B’s cis*-regulating roles and KDM2B’s function in promoting neuronal differentiation are conserved in human.

## Materials and methods Key resources table

### Contact for reagent and resource sharing

Further information and requests for resources and reagents should be directed to and will be fulfilled by the Lead Contact, Yan Zhou.

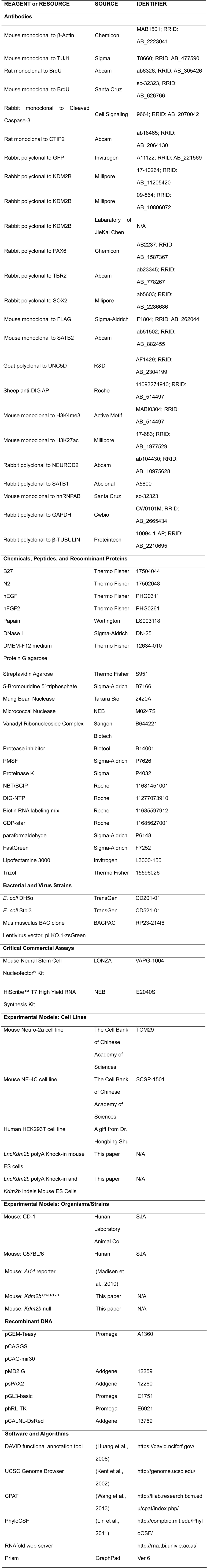
Key resources table.

### Experimental model and subject details

#### Mouse

All animal procedures were approved by the Animal Care and Ethical Committee of College of Life Sciences at Wuhan University. CD-1 and C57BL/6 mice were purchased from HNSJA. Mice were housed in a certified specific-pathogen-free (SPF) facility. The noon of the day on which the vaginal plug is found is counted as embryonic (E) day 0.5.

### Generation of *Kdm2b* ^*CreERT2/+*^ Knock-In Reporter Mice

*Kdm2b ^CreERT2/+^* knock-in reporter mice were generated by Biocytogen (Beijing, China). A sequence encoding the self-cleaving T2A peptide was fused in frame with exon 3 of the *Kdm2b* followed by the CreERT2-IRES-EGFP cassette. To generate the *Kdm2b* targeting vector, a 1 kb 5’ homology (LR), a 1 kb 3’ homology arm (RR), F2a-iCreERT2, or IRES-EGFP were amplified by PCR. Fragment LR and F2a-iCreERT2 were overlapped to form fragment LR-F2a-iCreERT2 (Sal I to BamH I). Fragment IRES-EGFP and RR were overlapped to form IRES-EGFP-RR (BamH I to Sac I). Then fragment LR-F2a-iCreERT2 and IRES-EGFP-RR were cloned into the TV-2G vector. For cloning the sgRNA-expression cassette, annealed DNA was ligated with pT7-sgRNA. SgRNAs were transcribed *in vitro* by T7 RNA Synthesis Kit (NEB). Targeting vector, Cas9 vector, and sgRNAs were microinjected into mouse zygotes. After injection, zygotes were immediately transferred into pseudo-pregnant female mice to generate founders, which were genotyped by PCR and sequencing. Positively founders were crossed with C57BL/6 wild-type mice to generate F1 mice. F1 mice were screened by PCR, and positive mice were confirmed by Southern blot using the iCre internal probe and 3’ external probe. The genders of embryos were not determined for analyses conducted in this study. See Supplementary file 1 - Table 4 for sgRNA sequences and genotyping primers.

### Generation of *Kdm2b* Null Mice

*Kdm2b* null mice were generated by Dr. Hongliang Li (Wuhan University, China). For cloning the sgRNA-expression cassette, sgRNAs targeting the exon encoding the CxxC domain were designed and synthesized, and annealed DNA was ligated to pT7-sgRNA. SgRNAs were transcribed *in vitro* by T7 RNA Synthesis Kit (NEB). Cas9 vector and sgRNAs were microinjected into the mouse zygote. After injection, the zygotes were immediately transferred into pseudo-pregnant female mice to generate founders, which were genotyped by PCR and sequencing. Positively founders were bred and crossed with C57BL/6 mice to generate F1 mice, which were screened by PCR and sequencing. See Supplementary file 1 - Table 4 for sgRNA sequences and genotyping primers.

### Genetic Lineage-tracing

All animals used for analyses in Figure 1 and Figure 1 - figure supplement 1 were heterozygous for the Cre allele (*Kdm2b^CreERT2/+^*).In Figure 1 - figure supplement 2, data were generated by crossing *Kdm2b^CreERT2/+^* with *Ai14^fi/fi^* animals, both with congenic *C57BL/6J* backgrounds. Tamoxifen was dissolved in corn oil as previously described (Guo et al., 2013). To perform lineage-tracing analyses using the *Kdm2b^CreERT2/+^;Ai14* mice, tamoxifen was injected into pregnant dams at indicated stages with a concentration of 100 mg/kg body weight.

### Generation of *LncKdm2b* polyA Knock-In (mESCs*^LncKdm2b-pASI+^*) and *Kdm2b* indels Mouse ES Cells

*LncKdm2b* polyA knock-in mouse ES cells (mESCs^*LncKdm2b-pAS/+*^) were generated by Biocytogen (Beijing, China). The targeting vector contains two homology arms (1 kb each), the 3 × SV40 polyA signal sequence and a BGH polyA signal (a total of 4 × polyA signals), followed by an expression cassette of ΔTK and Neo flanked by two loxP sites. The targeting vector was electroporated into mouse ES cells with Cas9-expressing vectors and sgRNAs that target the genomic site 1.8 kb downstream of the *LncKdm2b* TSS. Out of 200 neomycin resistant clones, one heterozygous knock-in ESC clone was obtained through PCR and sequencing analyses. To generate mESCs^*LncKdm2b-pAS/+*^ with *Kdm2b* indels, sgRNAs that target the second exon of *Kdm2b* were electroporated into mESCs^LncKdm2b-pAS/^+. mESC clones with distinguishable indel mutations between two alleles were selected by PCR and sequencing analyses. See Supplementary file 1 - Table 4 for sgRNA sequences, genotyping and qPCR primers.

### Cell Lines

HEK293T cells were gifts from Dr. Hongbing Shu (Wuhan University). Neuro-2a cells and NE-4C cells were purchased from the Cell Bank of Chinese Academy of Sciences Cells were maintained in indicated culture media (DMEM or MEM) containing 10% fetal bovine serum (Life Technologies or Hyclone) and used within ten passages since arrival.

### Plasmids Construction

For constructing eukaryotic expression vectors, full-length mouse *Kdm2b* was PCR amplified from the pMXs-Kdm2b-Flag vector, a gift from Dr. Baoming Qin (Guangzhou Institutes of Biomedicine and Health, Chinese Academy of Sciences), then subcloned into the pCAGGS vector using EcoR I / Mlu I. *Kdm2b*-mJmjC, *Kdm2b*-ΔCxxC, *Kdm2b*-mPHD, *Kdm2b*-ΔFbox, *Kdm2b*-ΔLRR, and shRNA-resistant mutants were constructed using site-directed mutagenesis. Full-length *LncKdm2b* was PCR amplified from the cDNA of the E16.5 C57BL/6 embryonic cortex, and the PCR product was cloned into pCAGGS using EcoR I / Not I. The longest ORF of *LncKdm2b* was fused in frame with sequence encoding C-terminal 3 × Flag tag and cloned into the eukaryotic expression vector pcDNA3.1 using EcoR I /Hhe I. The CDS sequence of mouse hnRNPAB was PCR amplified from the cDNA of the E16.5 C57BL/6 embryonic cortex, was cloned into the eukaryotic expression vector pFLAG-N3 (a gift from Dr. ZhiYin Song, Wuhan University) using Xho I / EcoR I in frame with sequence encoding C-terminal 3 × Flag tag. pCALNL was a gift from Dr. Xiaoqun Wang (Institute of Biophysics, Chinese Academy of Sciences). Luciferase reporter vector was constructed according to the previous study (Li et al., 2017b). Briefly, the T5 Forward, T5 Reverse, T5-mini Forward, or T5-mini Reverse were PCR amplified from the genomic DNA of C57BL/6 mice and cloned into pGL3-Basic Vector using Mlu I and Xho I. *Kdm2b* promoter and *Kdm2b* promoter with the CArG box deletion were cloned into pGL3-Basic Vector using Sac I and Xho I. For constructing RNA tethering vectors, the LacZ sequence from pcDNA3-BoxB-LacZ was removed by Xho I and Xba I digestion, and *LncKdm2b* was amplified from the *pCAGGS-LncKdm2b* Vector and cloned into the same sites with Xho I and Xba I. 5 × UAS-TK-Luc, pcDNA3-Gal4-λN, and pcDNA3-BoxB-LacZ were gifts from Dr. Xiang Lv (CAMS & PUMC). For constructing KUS-d2EGFP or KUSR-d2EGFP, EGFP and ODC (422-461 aa) were amplified by PCR, overlapped to form fragment d2EGFP (EcoR I to Bgl II) (Corish and Tyler-Smith, 1999). Then d2EGFP was cloned into the pCAGGS vector using EcoR I and Bgl II. pCAGGS-d2EGFP was digested by Apa I, followed by Mung Bean Nuclease (Takara Bio) modification and removal of the CAG promoter by Sal I digestion. KUS or KUSR were PCR amplified from the genomic DNA of C57BL/6 mice and cloned into the same site with Xho I. For construction short-hairpin RNA (shRNA) vectors, the oligonucleotides for shRNA targeting *Kdm2b* or *LncKdm2b* were cloned into pLKO.1-zsGreen or pCAG-mir30 vectors using Age I / EcoR I or Xho I / EcoR I. A scramble shRNA plasmid was used as a negative control. Primer sequences for all constructs were listed in Supplementary file 1 - Table 4.

### Protein Expression and Purification for hnRNPAB

Plasmids expressing Flag-tagged hnRNPAB were transfected into HEK293T cells. Cells were harvested after 2 days to achieve optimal expression. 2 × 10^8^ HEK293T cells were resuspended in lysis buffer [20 mM Tris pH 8.0, 100 mM NaCl, 1 mM PMSF, protease inhibitor cocktail (Biotool)] followed by sonication with 30% power output (3 minutes, 0.5 seconds on, 0.5 seconds off) on ice. After centrifugation at 12,000 rpm for 10 minutes at 4°C, supernatants were incubated with 50 μL anti-Flag agarose beads (Biotool) for 2 hours at 4°C. The agarose beads were washed 4 × 5 minutes with TBS buffer, and bound protein was eluted with 200 ng/μL 3 × FLAG peptide (Sigma-Aldrich, F4799) in TBS buffer at 4°C for 30 minutes. the eluted sample was ultrafiltrated and concentrated with 0.5 mL Amicon Ultra-centrifugal filters (Millipore, UFC501024). The concentration of purified protein was determined using Bicinchoninic Acid Protein Assay Kit (Beyotime Biotechnology) and by Western blot.

### Lentivirus Production and Cell Infection

To obtain lentiviral particles, HEK293T cells (5 × 10^6^ cells in a 10-cm dish) were transiently transfected with 12 μg pLKO.1 shRNA constructs, 6 μg of psPAX2 and 6 μg pMD2.G. The supernatant containing lentivirus particles was harvested at 48 hours after transfection, and filtered through Millex-GP Filter Unit (0.22 μm pore size, Millipore). Viral particles were then stored at −80°C ultra-cold freezer until use. Cortical NPCs infected by lentivirus at a low viral titer. Knockdown efficiency was evaluated by RT-qPCR analysis three days post-infection.

### Antisense Oligonucleotide (ASO) Treatment

Phosphorothioate-modified antisense oligodeoxynucleotides (ASOs) were synthesized at BioSune (Shanghai, China), and transduced into Neuro-2a cells using Lipofectamine^®^ 3000 (Thermo Fisher Scientific) according to the manufacturer’s protocol at 100 nM. For transfecting primary NPCs, ASOs were introduced into tertiary cortical NPCs derived from E14.5 CD-1 mouse cortex by nucleofection (Lonza) according to the manufacturer’s protocol at 1 μM for 1 × 10^6^ cells. Optimal programs and solutions of the Lonza Cell Line Nucleofector Kit for the ASO delivery were tested. Total RNAs were collected for RT-qPCR analysis two days post-transfection. Total protein was collected for immunoblotting analysis four days post-transfection. See Supplementary file 1 - Table 4 for ASO sequences.

### Knockout of *Lnckdm2b* and T5 by CRISPR/Cas9

CRISPR/Cas9-mediated genomic knockout was performed essentially as described previously (Cheng et al., 2016). Briefly, annealed oligonucleotides for sgRNAs targeting T5 or *Lnckdm2b’s* exon2 were cloned into a PiggyBac-based vector (pPB-sgRNA-Cas9). pPB-sgRNA-Cas9 and the transposase-expressing vector were mixed in a 1:1 ratio and co-transfected into NE-4C cells using lipofectamine 3000 or electroporated into E13.5 CD-1 mouse embryonic cortices. NE-4C cells that stably express Cas9 and sgRNAs through transposon-mediated random insertion were selected by flow cytometry and maintained as mono-clones for two weeks. Individual NE-4C clones (23-26 clones) were picked, expanded and analyzed by PCR genotyping. E15.5 cortical cells were isolated from embryonic cortices two days after electroporation and maintained in neurosphere culture medium for a week. Cortical cells that stably express Cas9 and sgRNAs were selected by flow cytometry and an aliquot was subjected to genomic DNA or RNA isolation followed by PCR genotyping and qPCR. See Supplementary file 1 - Table 4 for sgRNA sequences, genotyping and qPCR primers.

### Northern Blot

Dorsal forebrain tissues were resected from E14.5 and E16.5 mouse embryos under dissecting microscopes. Total RNAs were extracted twice using Trizol (Thermo Fisher). The polyA+ RNA fractions were enriched using the NEBNext Poly(A) mRNA Magnetic Isolation Module (NEB). About 1 μg of polyA+ RNA from each sample was subjected to formaldehyde denaturing agarose electrophoresis followed by transferring to positively charged nylon membrane with 20× SSC buffer (3.0 M NaCl and 0.3 M sodium citrate, pH 7.0). Membrane was UV-cross-linked and incubated with DIG-labeled RNA probes (*LncKdm2b*, 217-1307 nt) generated by in *vitro* transcription with the DIG-RNA Labeling Mix (Roche). Hybridization was done overnight at 65°C in DIG Easy Hyb Hybridization solution (Roche). Membranes were stringently washed three times in wash buffer 1 (0.1× SSC and 0.1% SDS) for 15 minutes at 65°C, then rinsed in wash buffer 2 [0.1 M maleic acid, 0.15 M NaCl, 0.3% Tween 20 (pH 7.5)] and incubated in blocking reagent (Roche) for 1 hour at room temperature. Subsequently, membranes were incubated with a 60,000-fold dilution of anti-DIG-AP Fab fragment (Roche) in blocking reagent for 30 minutes at room temperature, washed three times in wash buffer 2 for 10 minutes at room temperature, and immersed in detection buffer [0.1 M TrisHCl, 0.1 M NaCl (pH 9.5)] for 5 min. Anti-DIG-AP was detected using CDP-star chemiluminescent substrate for alkaline phosphatase (Roche) and X-ray film exposure. See Supplementary file 1 - Table 4 for primers used in generating Northern Blot probes.

### *In situ* Hybridization (ISH)

To make ISH probes, the 5’-overhang of forward primer was modified with a T7 promoter (See Supplementary file 1 - Table 4 for the primers used in ISH probes). Digoxigenin labeled riboprobes were transcribed using the DIG-RNA Labeling Mix (Roche). *In situ* Hybridization was performed as described (Li et al., 2017b). In brief, all solutions were prepared properly to avoid RNase contamination. Digoxigenin-labeled *LncKdm2b* and *Kdm2b* riboprobes were transcribed *in vitro* using NTP mix containing digoxigenin-labeled UTP (Roche). E12.5 CD-1 mouse embryos and E16.5 mouse brains were fixed in chilled 4% paraformaldehyde (Sigma) in 1 × PBS overnight followed by treatment of 20% sucrose in 1 × PBS overnight. Tissues were embedded in OCT, and 14 μm sections were cut onto slides using a Leica CM1950 cryostat. Sections were permeabilized with 2 μg/mL proteinase K (Sigma) for 10 minutes followed by acetylation in 0.1 M TEA (triethanolamine) solutions (10 mL 1 M TEA solution and 250 μL acetic anhydride in 90 mL DEPC treated ddH2O) for 10 minutes. Slides were blocked in hybridization buffer (50% deionized formamide; 5 × SSC, 5 × Denharts; 250 μg/mL yeast RNA; and 500 μg/mL herring sperm DNA) at room temperature (R/T) for 3 hours followed by incubating with 0.1-0.2 ng/μL digoxigenin-labeled riboprobe in hybridization buffer overnight at 60°C in humidified boxes. Slides were washed with 65°C 0.1 × SSC for three times (20 minutes each) followed by blocking with 10% heat-inactivated sheep serum in buffer B1 (0.1 M Tris-HCl, pH 7.4; 150 mM NaCl) at room temperature for 1 hour. Sections were incubated with an alkaline phosphatase-conjugated anti-digoxigenin antibody (1:5000, Roche) overnight at 4°C. After washing three times in buffer B1, sections were equilibrated twice in buffer B3 (0.1 M Tris-HCl; 0.1 M NaCl; 50 mM MgCl_2_; 0.1% Tween-20, pH 9.5) for 10 minutes. Colorization was performed using NBT/BCIP (Roche) containing B3 solutions at R/T overnight in the dark. Slides were dehydrated, cleared and mounted using gradient ethanol, xylene, and neutral balsam sequentially.Images were collected using a Nikon 80i microscope equipped with Nikon DS-FI1C-U3 camera system.

### Immunofluorescence (IF) and Immunoblotting

IF and immunoblotting were performed as described (Li et al., 2017b). For immunofluorescent staining, 4% paraformaldehyde (PFA) fixed 14 μm sections or cells were permeabilized and blocked with blocking buffer (3% heat-inactivated normal goat serum, 0.1% bovine serum albumin and 0.1% Triton-X 100 in 10 mM Tris-HCl, pH 7.4; 100 mM NaCl) for one hour at R/T. Sections were then incubated with primary antibodies diluted in blocking buffer overnight at 4°C or R/T. The next day, slides were washed three times for 10 minutes with 1 × PBS and incubated with second antibodies in blocking buffer at R/T for an hour. Slides were mounted with anti-fade solution with DAPI after PBS wash. For triple IF labeling of EGFP/PAX6/BrdU, sections were stained for EGFP/PAX6 antibodies first, then treated with 20 μg/mL proteinase K (Sigma) for 5 minutes followed by 2 mol/L HCl for 30 minutes before BrdU staining. All immunofluorescence comparing expression levels were acquired at equal exposure times. Immunoblotting assays were carried out according to standard procedures.

### RNA-seq Transcriptome Analysis

Dorsal forebrain (cortex) tissues were resected from E10.5 or E12.5 mouse embryos under dissecting microscopes. Total RNAs were extracted twice using Trizol (Thermo Fisher) and were treated with DNase I (NEB Biolabs). The integrity of RNAs was analyzed using Agilent Bioanalyzer 2100. Removal of ribosomal RNAs (rRNAs) and construction of libraries for standard strand-specific RNA-seq were performed using Illumina HiSeq 2000 in BGI Tech. Quality control reads alignment, and gene-expression analysis were also carried out in BGI Tech. Some low-quality RNA reads were present in original data. Thus, four kinds of reads were removed before mapping to the mouse genome. 1) Adaptor sequences; 2) Poor quality reads that Q5 or less mass value bases account for more than 50% of the entire reads; 3) Reads that have a proportion of “N” greater than 10%. 4) Reads that align with mouse rRNA. Next, the resulting clean reads were mapped to mouse genome (NCBI37/mm9) by TopHat (Trapnell et al., 2009) and an *ab initio* transcriptome reconstruction approach was performed by Cufflinks (Trapnell et al., 2012). To explore the expression patterns of coding and non-coding gene across embryonic cortical development, we used the Galaxy platform (Goecks et al., 2010) to integrate RNA-seq data from four other studies (mESCs and NPCs: GSE20851; mouse E14.5 VZ, IZ, and CP: GSE30765; E17.5 cortex: GSE39866; adult mouse cortex: GSE39866, GSE45282) (Ayoub et al., 2011; Dillman et al., 2013; Guttman et al., 2010; Ramos et al., 2013). Finally, we used Cuffnorm to calculate FPKM (Fragments Per Kilobase of exon per Million fragments mapped). GO analysis was performed using the DAVID Functional Annotation Bioinformatics Microarray Analysis tool (Huang et al., 2008). The RNA-seq data of E10.5 or E12.5 mouse cortice were deposited in the Gene Expression Omnibus with accession no. GSE55600.

### RNA Fractionation

RNA fractionation was performed as previously described (Cabianca et al., 2012). In brief, neural progenitor cells from E14.5 mouse cortices were detached by treating with 1 × Trypsin, counted and centrifuged at 168 g for 5 minutes. The pellet was lysed with 175 μL/10^6^ cells of cold RLN1 solution [50 mM Tris-HCl, pH 8.0; 140 mM NaCl; 1.5 mM MgCh; 0.5% NP-40; 2 mM Vanadyl Ribonucleoside Complex (Sangon Biotech)] for 5 minutes. The suspension was centrifuged at 4°C and 300 g for 2 minutes. The supernatant, corresponding to the cytoplasmic fraction, was transferred into a new tube and stored on ice. The pellet containing nuclei was corresponding to nuclear fractions.

Total RNA was extracted from the cytoplasmic and nuclear fractions using TRIzol solution. The samples were treated with DNase I, washed with 75% ethanol and then resolved in 30 μL RNase-free water. 1 μg of RNA was used for the first-strand synthesis with the PrimerScript™ Reverse Transcriptase (Takara Bio) using oligo-dT and random primers. cDNA was used for qPCR with iTaq^TM^ Universal SYBR^®^ Green Supermix (Biorad) and analyzed by a CFX Connect™ Real-Time PCR Detection System (Bio-rad). See Supplementary file 1 - Table 4 for qPCR primers.

### Fluorescent RNA *in situ* Hybridization (FISH) and Immunofluorescence Microscopy

FISH probes were designed using the Stellaris Probe Designer (Biosearch Technologies) (See Supplementary file 1 - Table 4 for primers used in FISH probes). A total of 38 probes with 20 nucleotides in length were used (Tsingke Biotech). Probes were biotinylated using terminal transferase (NEB, M0315S) with Bio-N6-ddATP (ENZO, ENZ-42809) as substrates. To detect *LncKdm2b* RNA, adherently cultured primary NPCs derived from E13.5 mouse cortex were rinsed in PBS and then fixed with 3.7% formaldehyde in PBS at room temperature for 10 minutes. Cells were permeabilized with 70% ethanol at 4°C overnight. Cells were treated with RNase A (100 μg/mL) or with PBS (in the control group) at 37°C for 1 hour. After washing in Wash Buffer A (Biosearch Technologies, SMF-WA1-60) for 5 minutes, cells were incubated with DNA probes in hybridization buffer (Biosearch Technologies, SMF-HB1-10) at 37°C overnight. After hybridization, cells were incubated with Alexa Fluor 555 conjugated streptavidin (1:1500 diluted in 1% BSA in PBS) at RT for 1 hour. Cells were washed twice with Wash Buffer A at 37°C for 30 minutes followed by nuclear counterstaining with DAPI. For colocalization studies, cells were co-stained with mouse anti-hnRNPAB (Santa Cruz Biotechnology).

### Luciferase Reporter Assays

To perform luciferase assays, Neuro-2a and HEK293T cells at ~60% confluency in each well of 24-well plates were transfected with 500 ng of pGL3-T5 Forward, pGL3-T5 Reverse, pGL3-T5-mini Forward, or pGL3-T5-mini Reverse plus 5 ng of pTK-Ren vectors using Lipofectamine^®^ 3000 (Thermo Fisher Scientific). For the RNA tethering experiment, Neuro-2a cells were grown in 24-well plates until 60% confluent and transfected with 150 ng of 5 × UAS-TK-Luc, pcDNA3-Gal4-λN, and pcDNA3-BoxB-LacZ or pcDNA3-BoxB-LncKdm2b, plus 5 ng of pTK-Ren vectors. For the detection of dose-dependent repression effect of *BoxB-LncKdm2b* on reporter activity, different doses of the pcDNA3-BoxB-LncKdm2b plasmid at 50 ng, 100 ng or 200 ng were used. As the amount of BoxB-lncRNA plasmid was increased, an equal amount of BoxB-LacZ was reduced accordingly. Twenty-four hours after transfection, cells were harvested and assayed for reporter activity using the Dual-Glo Luciferase Assay System and the GloMax multidectection system according to manufacturer’s instructions (Promega). Each data point was taken as the average Luc/Ren ratio of triplicate wells. To test the effects of hnRNPAB knockdown on *Kdm2b’s* promoter activity (CArG Box-containing), Neuro-2a cells at ~50% confluency in each well of 24-well plates were transfected with 50 nM siRNA targeting hnRNPAB. The second transfections with 500 ng pGL3-pKdm2b or pGL3-pKdm2b Δ CArG plus 5 ng of pTK-Ren vectors were done twenty-four hours later. Cells were harvested 24 hours later and assayed for reporter activity.

### Nuclear Run-on (NRO)

Nuclear run-on was performed as previously described (Roberts et al., 2015). About 1-4 × 10^6^ Neuro-2a cells were harvested and washed with PBS for one run-on experiment. Cell pellets were added 1 mL of lysis buffer (10 mM Tris-HCl, pH 7.4; 10 mM NaCl; 3 mM MgCh; 0.5% NP-40) and incubated on ice for 5 minutes. After centrifugation at 300 g for 4 minutes at 4°C, the pellet was washed with lysis buffer without NP-40 and resuspended with 40 μL nuclear storage buffer (50 mM Tris-HCl, pH 8.3; 40% glycerol; 5 mM MgCl_2_; 0.1 mM EDTA). Equal volume of 2 × transcription buffer [20 mM Tris-HCl, pH 8.3; 300 mM KCl; 5 mM MgCh; 4 mM DTT; 1 mM each of ATP, GTP and CTP, 0.5 mM UTP, 100 U RNase Inhibitor (Takara Bio)] was added into nuclei and then supplied with 0.5 mM BrUTP (Sigma). After incubation at 30°C for 30 minutes, RNA was extracted by TRIzol, and digested by 6 U DNase I (Thermo Fisher). About 30 μL of protein G agarose beads were washed with PBST, resuspended in 30 μL PBST. 2 μg of anti-BrdU monoclonal antibody (Santa Cruz) was added and incubated on a rotating platform for 10 minutes at room temperature. 150 μL of blocking buffer (0.1% PVP and 0.1% BSA in PBST) was added and incubated for 30 minutes at room temperature. After four times wash with 300 μL PBSTR (80 U RNase Inhibitor per 10 mL of PBST), pellets were resuspended in 100 μL of PBSTR. NRO-RNA samples were treated at 65°C for 5 minutes to denature RNA secondary structures and then incubated with the BrdU antibody-bound agarose for 60 minutes at room temperature on a rotating platform. After four times wash with 300 μL PBSTR, RNA was extracted by resuspending the pellet in 500 μL TRIzol. RNAs were reverse-transcribed, and detected by qPCR. RT-qPCR primers used to detect the pre-mRNAs of *LncKdm2b*, or *Kdm2b* were designed to cover one exon-intron junction, that is, one primer locates in the intron and the other in the adjacent exon. See Supplementary file 1 - Table 4 for qPCR primers.

### Chromatin immunoprecipitation (ChIP)

ChIP experiments were performed essentially as described previously (Wu et al., 2008). Briefly, 1 × 10^7^ Neuro-2a cells per experiment were crosslinked with 1% formaldehyde in the medium for 10 minutes at room temperature and quenched by adding 0.125 M glycine for 5 minutes. Cells were then washed twice with ice-cold PBS. Cells were then harvested in 500 μL Digestion buffer (50 mM Tris-HCl, pH 7.9; 5 mM CaCh; 100 μg/mL BSA) plus 1400 U Micrococcal Nuclease (NEB) for 20 minutes at 37°C, followed by adding 5 μL 0.5 M EDTA and incubating on ice for 5 minutes. Sonicate samples in EP tubes on ice with power output 30%, 3 minutes, 0.5 seconds on, 0.5 seconds off. One percent of the sonicated lysate was used as the input. Sonicated lysates were diluted into 0.1% SDS using dilution buffer (50 mM Tris-HCl, pH 7.6; 1 mM CaCh; 0.2% Triton X-100; 0.5% SDS) and incubated with 25 μL pre-washed protein G agarose beads plus 4 μg anti-hnRNPAB (Santa Cruz Biotechnology) or 2 μg anti-H3K4me3 (Active Motif) or anti-H3K27ac (Millipore) antibodies on rocker at 4°C overnight. After wash with Wash Buffer I (20 mM Tris-HCl, pH 8.0; 150 mM NaCl; 2 mM EDTA; 1% Triton X-100; 0.1% SDS), 4 times wash with Wash Buffer II (20 mM Tris-HCl, pH 8.0; 500 mM NaCl; 2 mM EDTA; 1% Triton X-100; 0.1% SDS), 4 times wash with Wash Buffer III (10 mM Tris-HCl, pH 8.0; 0.25 M LiCl; 1 mM EDTA; 1% deoxycholate; 1% NP-40), and 2 times wash with TE Buffer, resuspend each pellet in 100 μL elution buffer (0.1 M NaHCO3; 1% SDS) with 1 μL 20 mg/mL proteinase K. Cross-linked chromatin was reversed at 65°C overnight. DNAs were purified using the PCR purification Kit (TianGen). Purified DNA was used for quantitative PCR analyses and was normalized to input chromatin. See Supplementary file 1 - Table 4 for qPCR primers.

### Chromosome Conformation Capture (3C)

3C experiments were performed essentially as described previously (Hagege et al., 2007). Briefly, 1 × 10^7^ Neuro-2a cells per experiment were crosslinked with 2% formaldehyde in the medium for 10 minutes at room temperature and quenched by 0.125 M glycine for 5 minutes. Cells were then washed twice with ice-cold PBS. Cells were lysed in 3C lysis buffer (10 mM Tris-HCl, pH 8.0; 10 mM NaCl; 0.2% NP40; PMSF) for 1 hour on rocker at 4°C and nuclei were pelleted by centrifugation at 3500 rpm for 10 minutes at 4°C. Pellets were then resuspended in 500 μL 1.2 × restriction enzyme buffer (NEB) with 0.3% SDS and incubated with rotation at 37°C for 1 hour. Triton X-100 was added to a final concentration of 2% followed by 1 hour incubation at 37°C with rotation. 400 U of highly concentrated EcoR I (NEB) was added and incubated overnight at 37°C with rotation. The following day, SDS was added to a final concentration of 1.6%, and samples were incubated at 65°C for 20 minutes. Samples were then brought up to a final volume of 7 mL in T4 ligase buffer (66 mM Tris-HCl, pH7.6; 6.6 mM MgCl_2_; 10 mM DTT; 100 jM ATP) with 1% Triton X-100. Samples were rotated at 37°C for 1 hour. Samples were then chilled on ice for 5 minutes, and 700 U T4 DNA ligase (Takara Bio) was added, and samples were incubated at 16°C for overnight followed by 30 minutes at room temperature. Next, 300 μg of proteinase K was added, and crosslinks were reversed at 65°C overnight. The following day, an additional 300 μg of RNase A was added and incubated at 37°C for 40 minutes. Finally, genomic DNA was purified by phenol-chloroform extraction followed by ethanol precipitation. Ligation events were detected using specific primers. qPCRs were performed on a CFX Connect™ Real-Time PCR Detection System using iTaq^TM^ Universal SYBR^®^ Green Supermix (Bio-rad). Specificity and efficiency of all 3C primers were verified by performing digestion and ligation of the BAC DNA containing the regions of interest. Ligation products were then serially diluted in sheared genomic DNA, and the efficiency of each PCR reaction was verified. Amplicons from BAC qPCRs and actual 3C template were run on agarose gel to verify the production of a single band of the expected size. See Supplementary file 1 - Table 4 for qPCR primers.

### Biotin-labeled RNA pull-down

RNA pull-down was performed as described previously (Tsai et al., 2010). To make biotinylated RNA pull-down probes, the 5’-overhang of forward primer was modified with a T7 promoter (See Supplementary file 1 - Table 4 for the primers used in RNA pull-down probes). Biotinylated RNAs were transcribed using the Biotin-RNA Labeling Mix (Roche) and HiScribe™ T7 High Yield RNA Synthesis Kit (NEB) according to the manufacturer’s protocol. About 3 μg of biotinylated RNA was heated at 90°C for 2 minutes, and then cooled down on ice for 2 minutes in RNA structure buffer (10 mM Tris pH 7, 0.1 M KCl, 10 mM MgCl_2_), and then shifted to room temperature (RT) for 20 minutes. About 5 x 10^7^ primary cells from E14.5 mouse cortices were used for each RNA pull-down experiment. Cells were resuspended in 2 mL PBS, 2 mL nuclear isolation buffer (1.28 M sucrose; 40 mM Tris-HCl pH 7.5; 20 mM MgCl_2_; 4% Triton X-100), and 6 mL water for 20 minutes on ice. Nuclei were pelleted by centrifugation at 2,500 g for 15 minutes and resuspended in 1mL RIP buffer [150 mM KCl, 25 mM Tris pH 7.4, 0.5 mM DTT, 0.5% NP-40, 1 mM PMSF and protease Inhibitor cocktail (Biotool). Resuspended nuclei were sonicated on ice at 30% power output for 3 minutes (0.5 seconds on, 0.5 seconds off). Nuclear extracts were collected by centrifugation at 12,000 rpm for 10 minutes, and were pre-cleared by 40 μL Streptavidin agarose (Thermo Fisher) for 20 minutes at 4°C with rotation. Pre-cleared lysates were mixed with 3 μg folded biotinylated RNA and 20 μg yeast RNA at 4°C overnight, followed by adding 60 μL washed Streptavidin agarose beads to each binding reaction and incubating at RT for 1.5 hours. After 4 × 10 minutes washes by RIP buffer (containing 0.5% sodium deoxycholate) at 4°C, proteins bound to RNA were eluted in 1 x sample buffer by heating at 100°C for 10 minutes, and then subjected to SDS-PAGE, and further visualized by silver staining. Finally, proteins were identified by mass spectrometry.

### Native RNA-Protein Complex Immunoprecipitation

Native RNA-protein complex immunoprecipitation assays were carried out as described (Xing et al., 2017) with modifications. Dorsal forebrain (cortex) tissues were resected from E14.5 mouse embryos and homogenated in 1 mL lysis buffer [50 mM Tris pH 7.4, 150 mM NaCl, 0.5% NP-40, 1 mM PMSF, 2 mM RVC, protease inhibitor cocktail (Biotool)] followed by sonication with a 30% power output for 3 minutes (0.5 seconds on, 0.5 seconds off) on ice. After centrifuging at 12,000 rpm for 10 minutes at 4°C, the supernatant was pre-cleared with 30 μL protein G agarose beads. The pre-cleared lysates were further incubated with 4 μg anti-hnRNPAB antibody (Santa Cruz) for 2 hours at 4°C. Then 50 μL protein G agarose beads (blocked with 1% BSA and 20 μg/ml yeast tRNA) were added to the mixture and incubated for another 1 hours at 4°C followed by washing with wash buffer [50 mM Tris pH 7.4, 300 mM NaCl, 0.05% Sodium Deoxycholate, 0.5% NP-40, 1 mM PMSF, 2 mM RVC, protease inhibitor cocktail (Biotool)]. RNAs were extracted with Trizol. For qRT-PCR, each RNA sample was treated with DNase I (Thermo Fisher) and reverse transcription was performed with PrimerScript™ Reverse Transcriptase (Takara Bio) using random primers followed by qRT-PCR analysis. See Supplementary file 1 - Table 4 for qPCR primers.

### Formaldehyde Crosslinking RNA Immunoprecipitation

Formaldehyde crosslinking RNA immunoprecipitation assays were carried out as described (Xing et al., 2017) with modifications. Dorsal forebrain (cortex) tissues were resected from E14.5 mouse embryos and fixed 10 mL PBS with 1% formaldehyde 10 minutes at room temperature followed by incubation with 0.25 M glycine at room temperature for 5 minutes. After pelleting tissues at 1,000 rpm for 5 minutes, the pellet was homogenized in 1 mL RIPA buffer [50 mM Tris pH 7.4, 150 mM NaCl, 1% NP-40, 0.5% Sodium Deoxycholate, 1 mM PMSF, 2 mM RVC, protease inhibitor cocktail (Biotool)] followed by sonication with a 30% power output for 3 minutes (0.5 seconds on,0. 5 seconds off) on ice. After centrifuging at 12,000 rpm for 10 minutes at 4°C, the supernatant was pre-cleared with 30 μL protein G agarose beads and 20 μg/ml yeast tRNA at 4°C for 30 minutes. Then the pre-cleared lysate was incubated with 50 μL beads that were pre-coated with 4 μg anti-hnRNPAB antibody (Santa Cruz) for 4 hours at 4°C. The beads were washed 4 × 5 minutes with washing buffer I (50 mM Tris pH 7.4,I M NaCl, 1% NP-40, 1% Sodium Deoxycholate), and 4 × 5 minutes with washing buffer II (50 mM Tris pH 7.4, 1 M NaCl, 1% NP-40, 1% Sodium Deoxycholate, 1 M Urea). The RNA-protein complex was eluted from beads by adding 140 μL elution buffer (100 mM Tris pH8.0, 10 mM EDTA, 1% SDS) at room temperature for 5 minutes. To reverse crosslinking, 4 μL 5 M NaCl and 2 μL 10 mg/ml proteinase K were added into the RNA samples, and incubated at 42°C for 1 hours followed by incubation at 65°C for one hour. The RNA was extracted with Trizol. For qRT-PCR, each RNA sample was treated with DNase I (Thermo Fisher) and then reverse transcription was performed with PrimerScript™ Reverse Transcriptase (Takara Bio) using random primers followed by qRT-PCR analysis. See Supplementary file 1 - Table 4 for qPCR primers.

### RNA-protein in *vitro* binding experiments

RNA-protein in *vitro* binding experiments (Dig-RNA pull-down assays) were carried out as described (Xing et al., 2017) with modifications. To make Dig-labeled *LncKdm2b* truncations and loop1 mutations, the 5’-overhang of forward PCR primer was modified with a T7 promoter (See Supplementary file 1 - Table 4 for the primers used in RNA pulldown probes). Digoxigenin labeled riboprobes were transcribed using the DIG-RNA Labeling Mix (Roche). 2 × 10^7^ HEK293T cells (Two 10-cm dishes) expressing Flag-hnRNPAB were harvested and resuspended in 1 mL lysis buffer [50 mM Tris pH 7.4, 150 mM NaCl, 0.5% NP-40, 0.5 mM PMSF, 2 mM RVC, protease inhibitor cocktail (Roche)] followed by sonication on ice. After centrifuging at 12,000 rpm for 10 minutes at 4°C, the supernatant was incubated with 50 μL anti-Flag agarose beads (Biotool). After immunoprecipitation and washing 4 × 5 minutes with TBS buffer, one fifth beads was saved for immunoblotting. The rest was equilibrated in binding buffer [50 mM Tris-HCl at pH 7.4, 150 mM NaCl, 0.5% NP-40, 1 mM PMSF, 2 mM RVC, protease inhibitor cocktail (Biotool)] and incubated with 300 ng Dig-labeled RNA (Dig-labeled RNAs were annealed by heating at 65°C for 5 minutes followed with slowly cooling down to room temperature) for at 4°C for 4 hours. After washing 4 × 5 minutes with binding buffer, the bound RNA was extracted with Trizol and analyzed by Northern blotting.

### Electrophoretic Mobility Shift Assay (EMSA)

For RNA EMSAs, digoxigenin labeled riboprobes (*LncKdm2b* loop1, 463-625 nt) were transcribed using the DIG-RNA Labeling Mix (Roche). EMSA experiments were conducted according to the manufacturer’s protocol with a Light Shift Chemiluminescent RNA EMSA kit (Thermo Scientific). The Dig-labeled RNAs were annealed by heating at 65°C for 5 minutes followed with slowly cooling down to room temperature. One fmol labeled RNAs were used for each EMSA reaction. For detection of dose-dependent binding of protein to RNA, different doses of Flag-hnRNPAB (1 μg, 3 μg or 10 μg) were used. Unlabeled probe was used for competitive reaction. Binding reactions were incubated in binding buffer [10 mM HEPES pH7.5, 20 mM KCl, 1 mM MgCl2, 1 mM DTT] at room temperature for 25 minutes, then immediately loaded onto a 2% nondenaturing agarose 0.5 × TBE (Tris-borate-EDTA) gel. After transfer to a nylon membrane, labeled probes were cross-linked by UV, probed with Anti-Digoxigenin-AP antibody, and incubated with detection substrates.

### Real-Time Quantitative Reverse Transcription PCR (qRT-PCR)

Total RNAs (0.5 - 1 μg) were reverse-transcribed at 42°C using PrimerScript™ Reverse Transcriptase (Takara Bio). Then iTaq^TM^ Universal SYBR^®^ Green Supermix (Bio-rad) was employed to perform quantitative PCR on a CFX Connect™ Real-Time PCR Detection System (Bio-rad). Gene expressions were determined using the 2^-ΔΔCt^ method, normalizing to housekeeping genes *Gapdh*. See Supplementary file 1 - Table 4 for qPCR primers.

### *In Utero* Electroporation of Developing Cerebral Cortices

*In utero* microinjection and electroporation were performed essentially at E13.5 as described (Li et al., 2017b). In brief, pregnant CD-1 mice were anesthetized by intraperitoneal injection of pentobarbital (70 mg/kg). The uteri were exposed through a 2 cm midline abdominal incision. Embryos were carefully pulled out using ring forceps through the incision and placed on sterile and irrigated drape. Intermittently wet uterine walls with saline to prevent drying. Supercoiled plasmid DNA (prepared using Endo Free plasmid purification kit, Tiangen) mixed with 0.05% Fast Green (Sigma) was injected through the uterine wall into the telencephalic vesicle of 3-4 embryos at intervals using pulled borosilicate needles (WPI). Electric pulses (36 V, 50 ms duration at 1 s intervals for 5 times) were generated using CUY21VIVO-SQ (BEX) and delivered across the uterine wall using 5 mm forceps-like electrodes (BEX). The uteri were then carefully put back into the abdominal cavity and incisions were sutured. The whole procedure was completed within 30 minutes. Mice were warmed on a heating pad until they woke up and given analgesia treatment (Ibuprofen) in drinking water until sacrifice.

### Primary Culture of Embryonic Cortical Neural Progenitor Cells (NPCs)

Primary culture of embryonic cortical NPCs was performed as described (Li et al., 2017b). In brief, E11.5 or E12.5 mouse cortices (dorsal forebrain) tissues were washed with and minced in filter-sterilized hibernation buffer (30 mM KCl; 5 mM NaOH; 5 mM NaH2PO4; 5.5 mM glucose; 0.5 mM MgCl_2_; 20 mM Na-pyruvate; 200 mM Sorbitol, pH 7.4) followed by dissociating into single cells using pre-warmed Papain (Worthington Biochemical) enzyme solution (1 × DMEM; 1 mM Na-pyruvate; 1 mM L-Glutamine; 1 mM N-Acetyl-L-Cysteine; 20 U/mL Papain; 12 μg/mL DNase I). Dissociated cells were cultured using serum-free media consisting of DMEM/F12 media (Life Technologies), N2 and B27 supplements (1 ×, Life Technologies), 1mM Na-pyruvate, 1 mM N-Acetyl-L-Cysteine (NAC), human recombinant FGF2 and EGF (20 ng/mL each; Life Technologies). For adherent cortical cultures in Figure 2C and S3A-S3B, cells were maintained on poly-L-lysine coated plates with the presence of 20 ng/mL FGF2 for 24 hours followed by differentiation (FGF2 withdrawal) for 48 hours. For sphere culture in Figure 3F, cells were cultured with the presence of EGF and FGF2 for 1 week. For clonal culture in Figure 7A-7G, cells were maintained 72 hours with the presence of 20 ng/mL FGF2 for 72 hours (4 × 10^4^ cells per well in 24-well plates).

### Quantification and Statistical Analysis

Data were presented as the mean ± SEM unless otherwise indicated. Statistical analyses were conducted using GraphPad Prism (version 6.01). Statistical significance was determined using unpaired 2-tailed Student’s t test, 1-way ANOVA followed by Tukey post hoc test, 2-way ANOVA followed by the Bonferroni post hoc test, and linear regression when appropriate. *p*≤0.05 was considered statistically significant. ‘*’; *p* values less than 0.01, or 0.001 was marked as ‘**’, and ‘***’ respectively.

## Author Contributions

Conceptualization, W.L., W.S. and Y.Z.; Methodology, W.L. and W.S.; Investigation, W.L., W.S., B.Z., K.T., Y.L., L.M., Z.L, X.Z., and Y.L.; Writing – Original Draft, W.L. and Y.Z.; Writing – Review & Editing, W.L. and Y.Z.; Funding Acquisition, Y.L. and Y.Z.; Resources, X.W.; Supervision, Y.Z.

## Acknowledgements

We thank Drs. Hongliang Li, Haohong Li, Jiekai Chen, Baoming Qin, Xiaoqun Wang, Xiang Lv, Min Wu and Haining Du for sharing reagents and technical supports. We thank Drs. Weimin Zhong, Qin Shen and Xiaohua Shen, and all members in Zhou lab for critical reading of the manuscript. This work was supported by grants from National Natural Science Foundation of China (No. 31471361 and No. 31671418), National Key Basic Research Program of China (No. 2011CBA01102), National Natural Science Foundation of Hubei Province (2018CFA016) and Fundamental Research Funds for the Central Universities (2042017kf0205 and 2042017kf0242) to Yan Zhou.

---

**Figure 1 - figure supplement 1.**
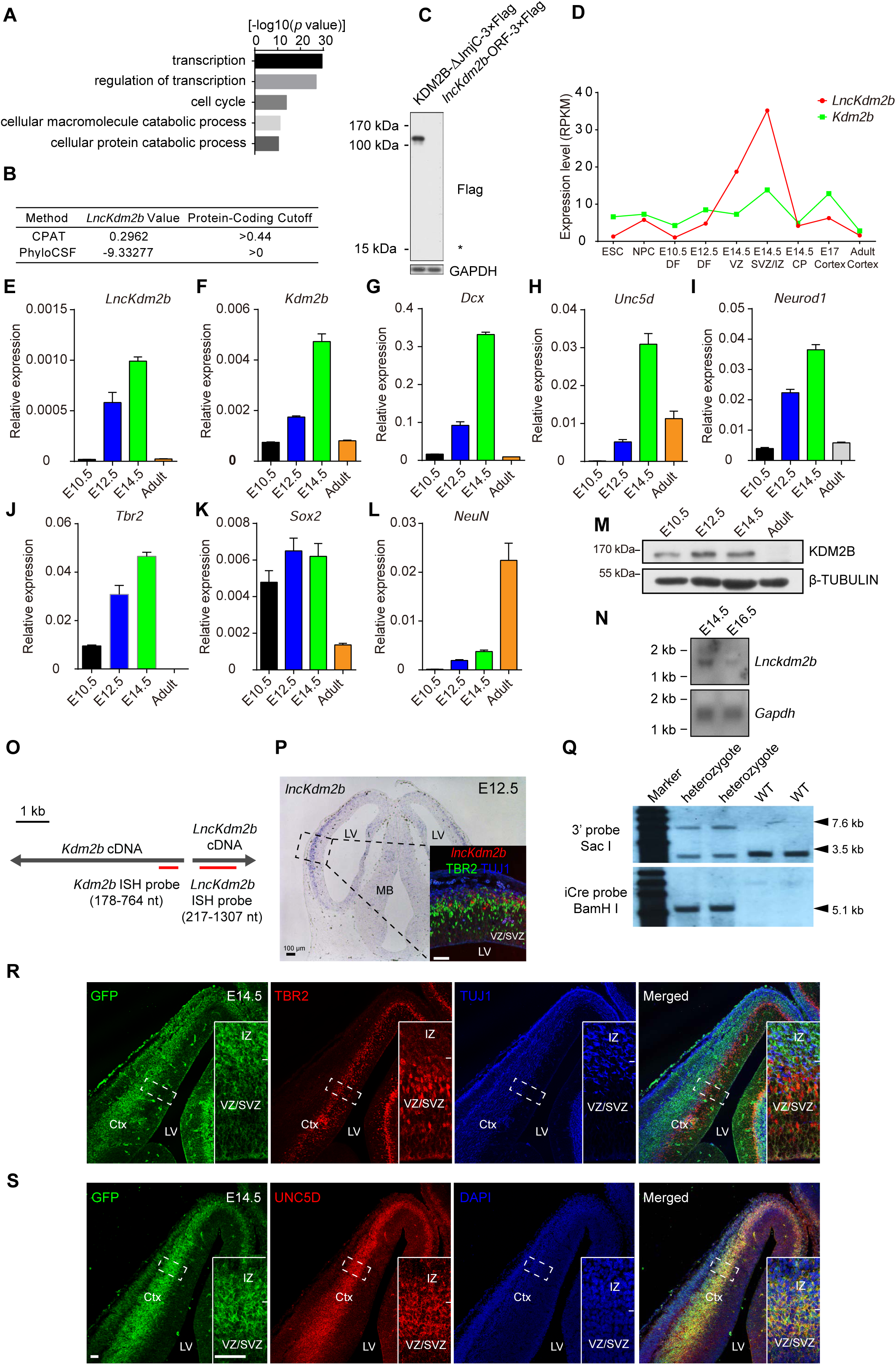
*LncKdm2b* and *Kdm2b* are Transiently Expressed in the Developing Mouse Embryonic Cortex. Related to Figure 1. (A) Gene ontology (GO) analysis of coding genes associated with divergent lncRNAs. The top GO terms (>1.5-fold and *P* < 1×10^-6^) are shown. (B) Protein-coding scores of *LncKdm2b* using CPAT and PhyloCSF programs. (C) *LncKdm2b*’s longest ORF fused with the 3 × Flag tag sequence at its 3’ was cloned into pcDNA3.1 vector for HEK293T cell transfection. After 48 hours, immunoblotting was performed to detect Flag-tagged proteins. *Kdm2b*-ΔJmjc tagged with 3 × Flag tag served as a positive control. Data are representative of three independent experiments. ‘*’ indicates predicted molecular weight encoded by *Flag-tagged LncKdm2b’s* ORF. (D) Fragments per kilobase per million mapped reads (FPKM) values for *LncKdm2b*, and *Kdm2b* in mouse ESCs, mouse NPCs and mouse cortices at indicated developmental stages. (E-L) RT-qPCR analyses of indicated markers of E10.5, E12.5, E14.5 and adult (6 weeks) mouse dorsal forebrains. The y-axis represents relative expression normalized to *Gapdh*. (M) Representative immunoblotting of mouse dorsal forebrains using antibodies against KDM2B and β-TUBULIN. (N) Northern blots of *LncKdm2b* and *Gapdh* using poly(A) RNAs extracted from mouse dorsal forebrains. (O) Schematic illustration of *in situ* hybridization probes for mouse *LncKdm2b* and *Kdm2b*. (P) *In situ* hybridization (ISH) of *LncKdm2b* on coronal sections of E12.5 mouse forebrain (the CD-1 strain). Scale bars, 100 μm. The higher-magnification image of the boxed area shows immunofluorescent staining for TBR2 (green) and TUJ1 on ISH section of *LncKdm2b* (red). Scale bars, 5O μm. (Q) Southern blot analysis of genomic DNA from wild-type (WT) or *Kdm2b^CreERT2/+^* knock-in mice. (R-S) Immunofluorescent staining for EGFP (green), TBR2 (red), TUJ1 (blue), UNC5D (red), and DAPI (blue) on cortical sections of E14.5 *Kdm2b^CreERT2/+^* mice. Boxed areas are enlarged at the bottom-right corners. Scale bars, 5O μm. In (E-L), quantification data are shown as mean + SD (n = 3 unless otherwise indicated). Ctx, cortex; LV, lateral ventricl; VZ, ventricular zone; SVZ, subventricular zone; IZ, intermediate zone.

**Figure 1 - figure supplement 2.**
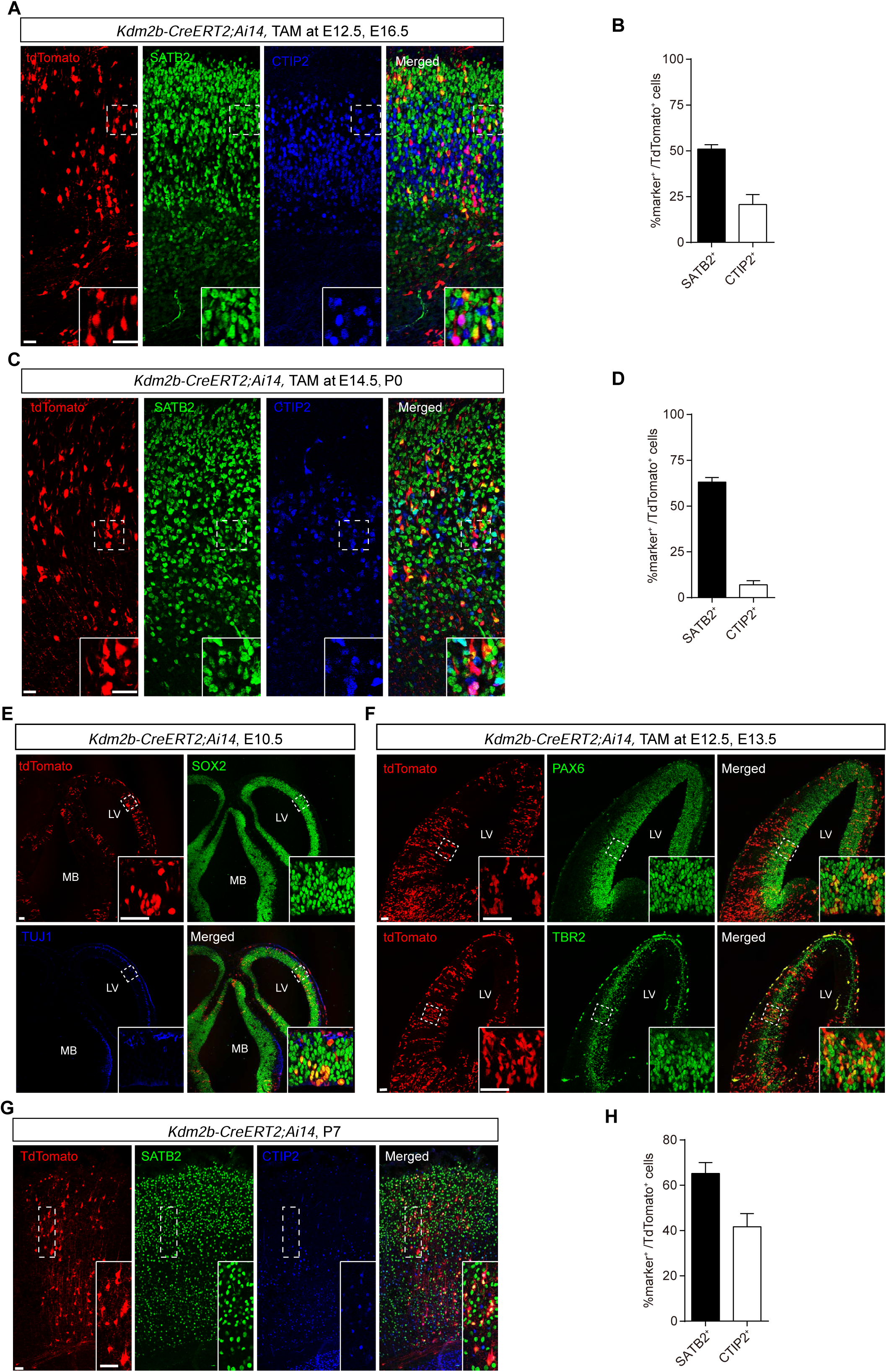
*Kdm2b*-expressing cortical cells are fated to be cortical projection neurons. Related to Figure 1. (A-D) Immunofluorescent staining for SATB2 and CTIP2 on coronal cortical sections of E16.5 (A) and PO (C) *Kdm2b^CreERT2/+^;Ai14* embryos. Tamoxifen (TAM, 1OO mg/kg) was injected at E12.5 or E14.5 respectively. Boxed areas are enlarged at the bottom-right corners. Quantification of SATB2 or CTIP2 expression in tdTomato+ recombined cells (B and D). A total of 3372 cells from 2 embryos were analyzed in (B), and 2282 cells from 2 animals in (D). (E) Immunofluorescent staining for SOX2 and TUJ1 on head sections of E1O.5 *Kdm2b^CreERT2/+^;Ai14* embryo. Boxed areas are enlarged at the bottom-right corners. (F) Immunofluorescent staining for PAX6 (top) and TBR2 (bottom) on head sections of E13.5 *Kdm2b^CreERT2/+^;Ai14* embryo. Tamoxifen (TAM, 1OO mg/kg) was injected at E12.5. Boxed areas are enlarged at the bottom-right corners. (G-H) Immunofluorescent staining for SATB2 and CTIP2 on coronal cortical sections of P7 *Kdm2b^CreERT2/+^;Ai14* mouse brain. Boxed areas are enlarged at the bottom-right corners.Quantification of SATB2 or CTIP2 expression in tdTomato+ recombined cells (H). A total of 373 cells were analyzed. In (B), (D), and (H), quantification data are shown as mean + SEM. LV, lateral ventricle; MB, midbrain. Scale bars, 50 μm.

**Figure 2 - figure supplement 1.**
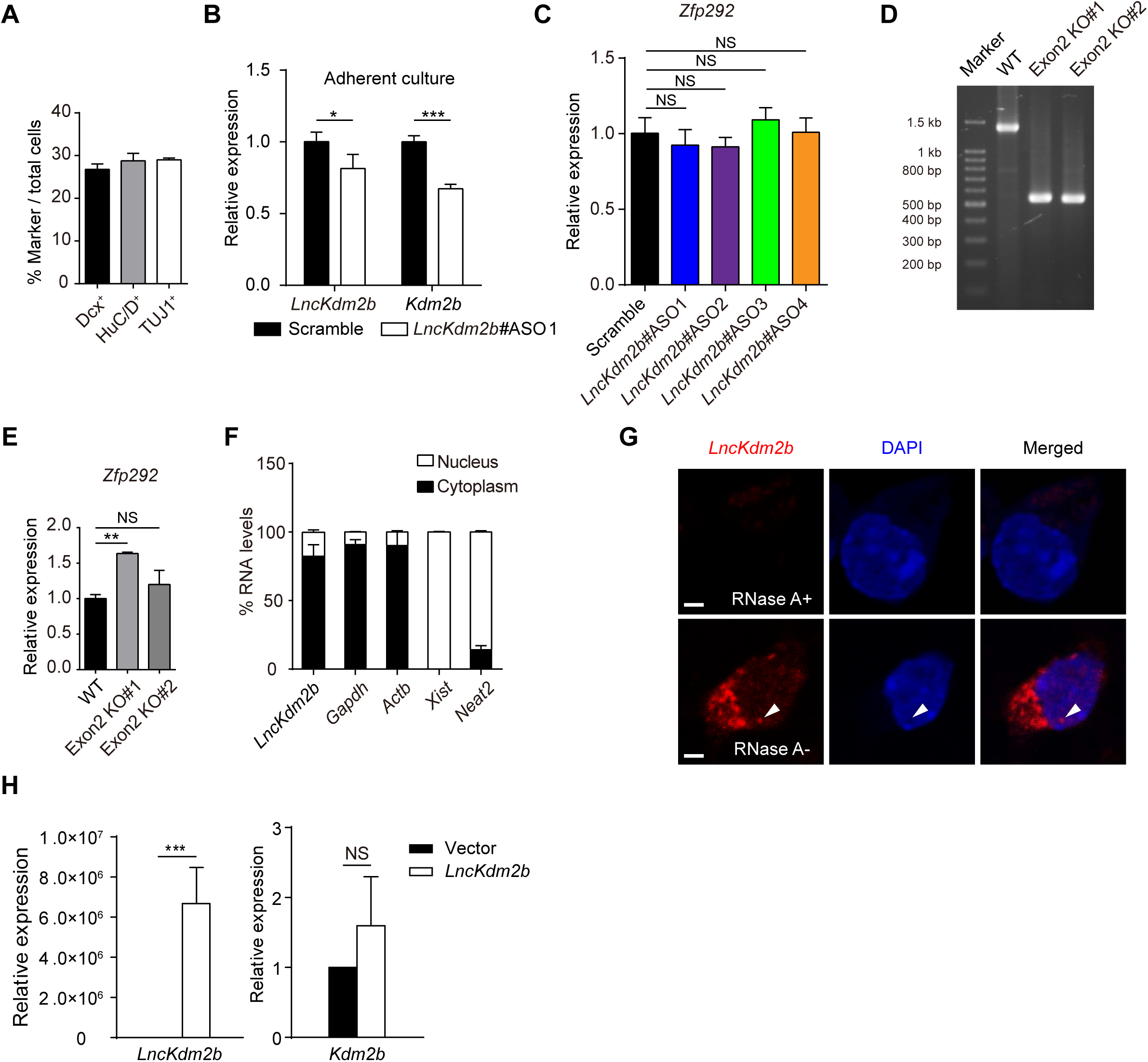
*LncKdm2b* maintains *Kdm2b* Transcription in *cis*. Related to Figure 2. (A) Percentages of marker-expressing neuronal cells in adherent cultures derived from E12.5 cortices. (B) RT-qPCR analysis of *LncKdm2b* and *Kdm2b* RNA levels in adherent cultures derived from E12.5 cortices. The cultures were treated with indicated ASOs. The y-axis represents relative expression normalized to *Gapdh*. (C) RT-qPCR analysis of *Zfp292* mRNA levels in Neuro-2a cells treated with indicated *LncKdm2b* ASOs for three days. The y-axis represents relative expression normalized to *Gapdh*. (D) Genotyping of NE-4C clones with *LncKdm2b’s* exon2 knocked out. (E) RT-qPCR analysis of *Zfp292* mRNA levels in NE-4C clones with *LncKdm2b’s* exon2 knocked out. The y-axis represents relative expression normalized to *Gapdh*. (F) RT-qPCR analysis for *LncKdm2b, Gapdh, Actb, Xist*, and *Neat2* from cytoplasmic and nuclear RNA fractions of primary E14.5 cortical neural progenitor cells (NPCs) cultured *in vitro* for four days. (G) Fluorescent *in situ* hybridization of *LncKdm2b* on cortical NPCs treated with or without RNase A. The nuclei were counter-stained with DAPI. (H) RT-qPCR analysis of *LncKdm2b* and *Kdm2b* mRNA levels in Neuro-2a cells transfected with empty or *LncKdm2b*-expressing vectors for 48 hours. The y-axis represents relative expression normalized to *Gapdh*. In (A), quantification data are shown as mean + SEM. In (B-C), (E-F), and (H) quantification data are shown as mean + SD (n = 3 unless otherwise indicated). In (B) and (H), statistical significance was determined using 2-tailed Student’s t test. In (C) and (E), statistical significance was determined using 1-way ANOVA followed by the Turkey’s post hoc test. * *p*<0.05, ** *p*<0.01, *** *p*<0.001, “NS” indicates no significance.

**Figure 3 - figure supplement 1.**
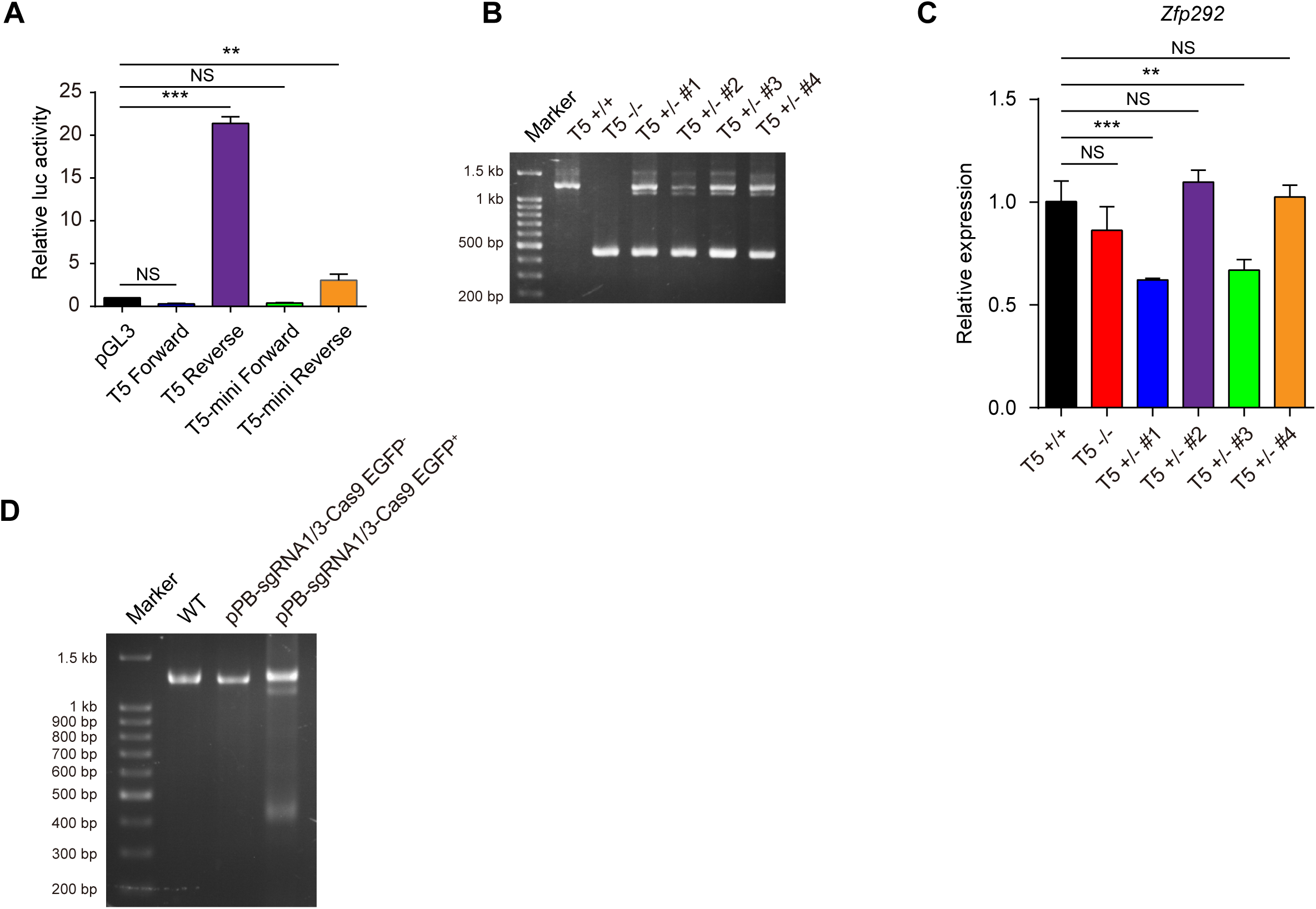
*LncKdm2b* maintains *Kdm2b* Transcription in *cis*. Related to Figure 3. (A) Luciferase activities in experiments where indicated vectors were transfected into HEK293T cells for 24 hours. ‘Forward’ and ‘Reverse’ indicate directions same as or opposite of *Kdm2b’s* transcription orientation. (B) Genotyping of NE-4C cells with the T5 region knocked out. (C) RT-qPCR analysis of *Zfp292* mRNA levels in NE-4C clones with the T5 region knocked out. The y-axis represents relative expression normalized to *Gapdh*. (D) Genotyping of cortical cells subjected to Cas9-mediated knockout of the T5 region. In (A) and (C), quantification data are shown as mean + SD (n = 3 unless otherwise indicated). In (A) and (C), statistical significance was determined using 1-way ANOVA followed by the Turkey’s post hoc test. * *p*<0.05, ** *p*<0.01, *** *p*<0.001, “NS” indicates no significance.

**Figure 4 - figure supplement 1.**
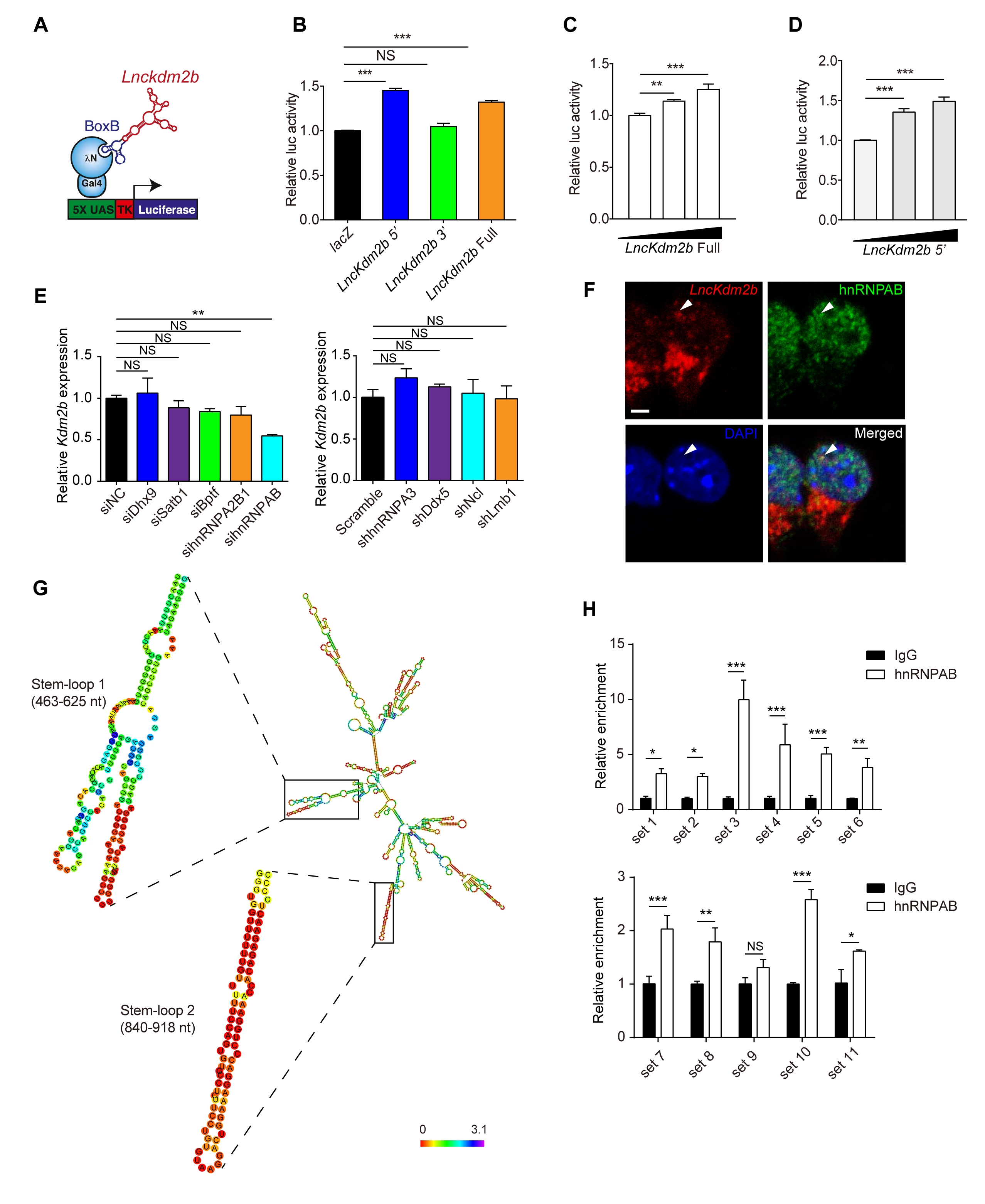
Characterization of *LncKdm2b*-Associated Proteins. Related to Figure 4. (A) The illustration describing the Gal4-λN/BoxB RNA tethering system. (B-D) Relative luciferase activities in experiments where Neuro-2a cells were transfected with plasmids expressing BoxB-tagged *LacZ*, full-length *LncKdm2b*, 5’ *LncKdm2b*, or 3’ *LncKdm2b* along with Gal4-λN and 5×UAS-TK-Luciferase-expressing plasmids for 24 hours. (E) RT-qPCR analysis of *Kdm2b* mRNA levels in Neuro-2a cells treated for three days with scramble siRNA (siNC) or siRNA targeting indicated molecules. The y-axis represents relative expression normalized to *Gapdh*. (F) Fluorescent *in situ* hybridization of *LncKdm2b* on cortical NPCs followed by co-staining of hnRNPAB and DAPI. (G) RNA secondary structure prediction by *RNAfold* showed two putative stem-loop structures. (H) ChIP-qPCR analysis of indicated primer sets enriched by anti-hnRNPAB antibodies in Neuro-2a cells. The y-axis shows fold enrichment normalized to the IgG control. In (B–E) and (H), quantification data are shown as mean + SD (n = 3 unless otherwise indicated). In (B–E), statistical significance was determined using 1-way ANOVA followed by the Turkey’s post hoc test. In (H), statistical significance was determined using 2-tailed Student’s t test.* *p*<0.05, ** *p*<0.01, *** *p*<0.001, “NS” indicates no significance.

**Figure 5 - figure supplement 1.**
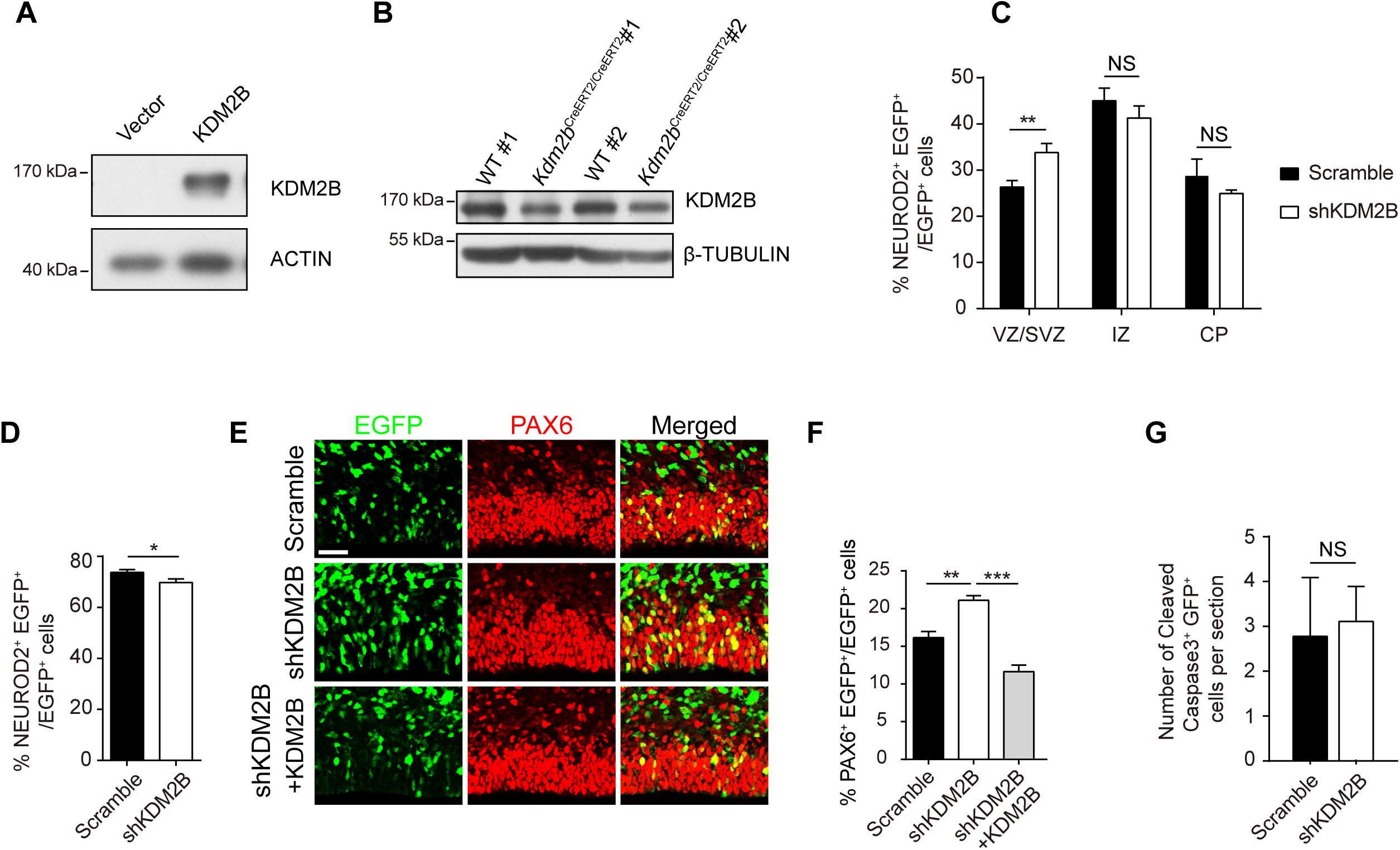
*Kdm2b* promotes Cortical Neurogenesis. Related to Figure 5. (A) Representative immunoblots of HEK293T cells transfected with empty vector or *Kdm2b*-expressing vector for two days using antibodies against KDM2B and ACTIN. (B) Immunoblotting of E15.5 embryonic cortices with indicated genotypes using antibodies against KDM2B and β-TUBULIN. (C-D) Quantification of relative location (C) and percentiles (D) of NEUROD2+ transduced cells in scramble or KDM2B shRNA electroporated sections. Three embryos each. (E-F) E13.5 mouse cortices were electroporated with indicated combination of vectors, with transduced cells labeled with EGFP. Embryos were sacrificed at E16.5 for immunofluorescent analysis. Representative VZ/SVZ images of sections immunostained with PAX6 (E) and quantification of PAX6+ transduced cells (F) were shown. Three embryos in control and shKDM2B, five embryos in shKDM2B plus KDM2B. Scale bars, 50 μm. (G) Quantification of Cleaved Caspase3+ transduced cells in scramble or KDM2B shRNA electroporated sections. Three embryos each. In (C-D) and (F-G), quantification data are shown as mean + SEM. In (C-D) and (G), statistical significance was determined using 2-tailed Student’s t test. In (F), statistical significance was determined using1-way ANOVA followed by the Turkey’s post hoc test. * *p*<0.05, ** *p*<0.01, *** *p*<0.001, “NS” indicates no significance. VZ, ventricular zone; SVZ, subventricular zone; IZ, intermediate zone; CP, cortical plate.

**Figure 6 - figure supplement 1.**
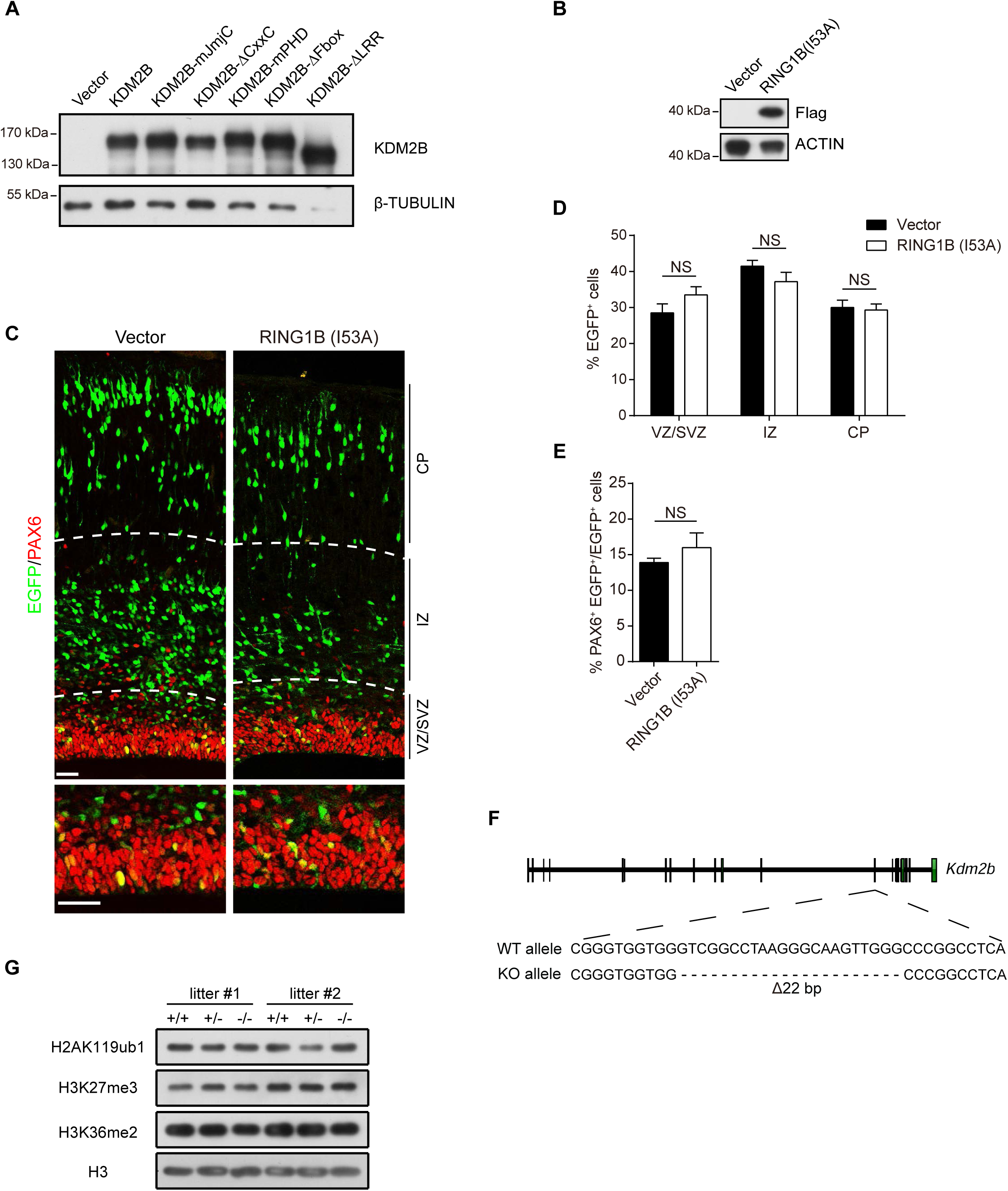
KDM2B promotes cortical neurogenesis independent of the Polycomb Repressive Complex 1 (PRC1). Related to Figure 6. (A) Representative immunoblotting of HEK293T cells transfected with indicated vectors for two days using antibodies against KDM2B and β-TUBULIN. (B) Representative immunoblotting of HEK293T cells transfected with empty and FLAG-RING1B(I53A)-expressing vectors for two days using antibodies against FLAG and ACTIN. (C-E) E13.5 mouse cortices were electroporated with empty or RING1 B(I53A)-expressing vector, along with EGFP-expressing vectors to label transduced cells. Embryos were sacrificed at E16.5 for PAX6 immunofluorescent analysis (C). The relative location of EGFP+ cells (D) and percentiles of PAX6+ transduced cells (E) were quantified. Six embryos in control, and four embryos in RING1B(I53A). Scale bars, 50 μm. (F) Schematic diagram showing generation of *Kdm2b* knockout mice using CRISPR/Cas9 mediated gene editing. (G) Representative immunoblotting of E9.5 mouse embryos with indicated genotypes using antibodies against H2AK119ub1, H3K27me3, H3K36me2, and H3. In (D–E), quantification data are shown as mean + SEM. In (D-E), statistical significance was determined using 2-tailed Student’s t test. * *p*<0.05, ** *p*<0.01, *** *p*<0.001, “NS” indicates no significance. VZ, ventricular zone; SVZ, subventricular zone; IZ, intermediate zone; CP, cortical plate.

**Figure 7 - figure supplement 1.**
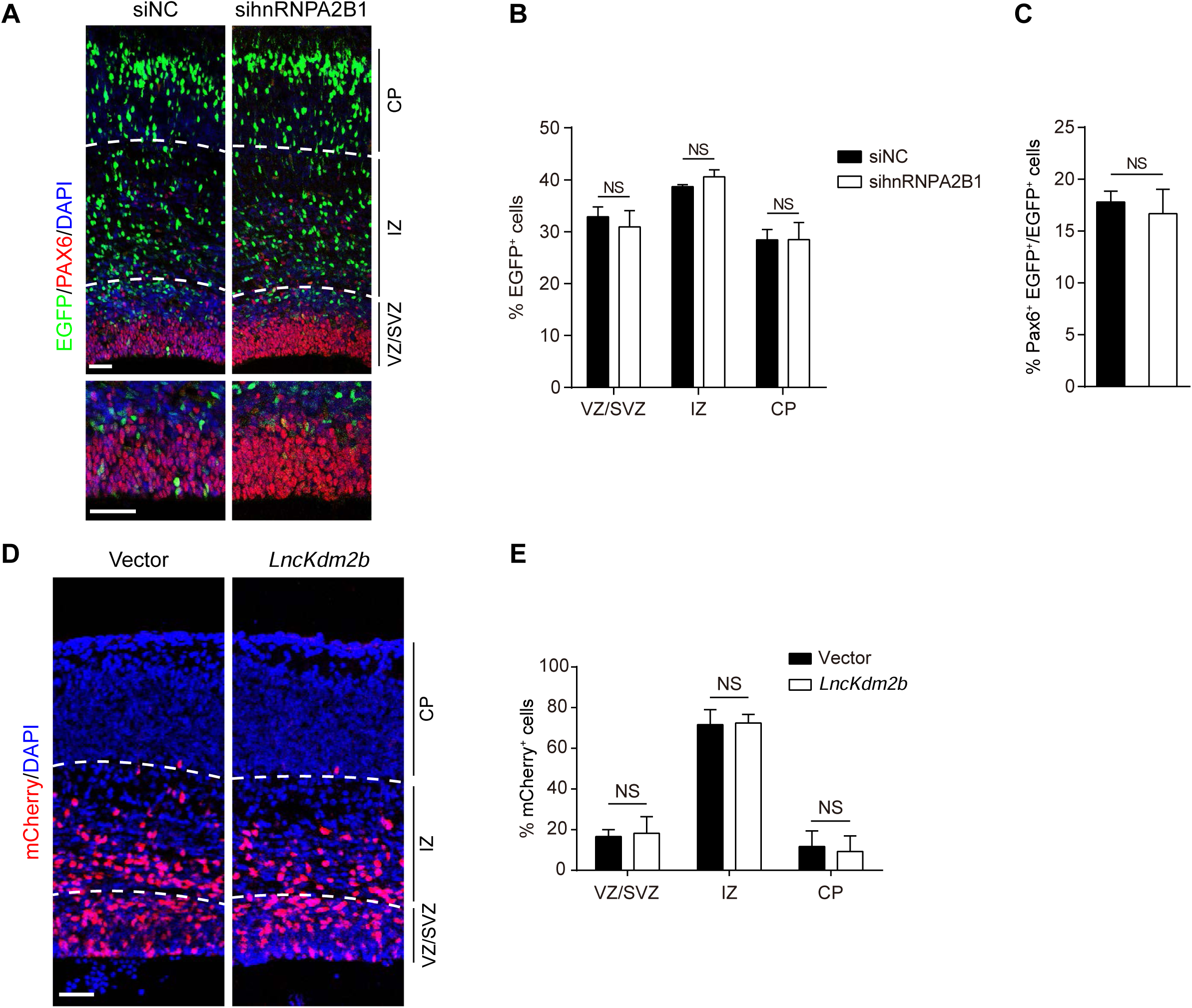
*LncKdm2b* Maintains Mouse Cortical Neurogenesis through KDM2B. Related to Figure 7. (A-C) E13.5 mouse cortices were electroporated with indicated siRNAs, with transduced cells labeled with EGFP. Embryos were sacrificed at E16.5 for PAX6 immunofluorescent stainings (A). The relative location of EGFP+ cells (B) and percentiles of PAX6+ transduced cells (C) were quantified. Three embryos in control (siNC), five embryos in sihnRNPA2B1. Scale bars, 50 μm. (D-E) E13.5 mouse cortices were electroporated with empty or *LncKdm2b*-expressing vector, along with mCherry-expressing vector to label transduced cells. Embryos were sacrificed at E15.5 followed by DAPI staining of coronal sections (D). The locations of mCherry+ cells were quantified (E). Seven embryos each. Scale bars, 50 μm. In (B–C) and (E), quantification data are shown as mean + SEM. In (B-E), statistical significance was determined using 2-tailed Student’s t test. * *p*<0.05, ** *p*<0.01, *** *p*<0.001, “NS” indicates no significance. VZ, ventricular zone; SVZ, subventricular zone; IZ, intermediate zone; CP, cortical plate.

## Additional files Supplementary file 1

Supplementary tables.

(1) Table 1. Divergent lncRNAs identified in this study.

(2) Table 2. Significantly-enriched proteins in *LncKdm2b*-precipitating extracts compared to the antisense-*LncKdm2b*.

(3) Table 3. Statistical analyses of electroporated cortices.

(4) Table 4. Sequences for all primers used in this study.

## References

Andricovich, J., Kai, Y., Peng, W., Foudi, A., and Tzatsos, A. (2016). Histone demethylase KDM2B regulates lineage commitment in normal and malignant hematopoiesis. The Journal of clinical investigation 126, 905–920.

Aprea, J., Prenninger, S., Dori, M., Ghosh, T., Monasor, L.S., Wessendorf, E., Zocher, S., Massalini, S., Alexopoulou, D., Lesche, M., et al.(2013). Transcriptome sequencing during mouse brain development identifies long non-coding RNAs functionally involved in neurogenic commitment. EMBO J 32, 3145–3160.

Arron, J.R., Winslow, M.M., Polleri, A., Chang, C.P., Wu, H., Gao, X., Neilson, J.R., Chen, L., Heit, J.J., Kim, S.K., et al.(2006). NFAT dysregulation by increased dosage of DSCR1 and DYRK1A on chromosome 21. Nature 441, 595–600.

Ayala, R., Shu, T., and Tsai, L.H. (2007). Trekking across the brain: the journey of neuronal migration. Cell 128, 29–43.

Ayoub, A.E., Oh, S., Xie, Y., Leng, J., Cotney, J., Dominguez, M.H., Noonan, J.P., and Rakic, P. (2011). Transcriptional programs in transient embryonic zones of the cerebral cortex defined by high-resolution mRNA sequencing. Proceedings of the National Academy of Sciences 108, 14950–14955.

Baek, Seung T., Kerjan, G., Bielas, Stephanie L., Lee, Ji E., Fenstermaker, Ali G., Novarino, G., and Gleeson, Joseph G. (2014). Off-Target Effect of doublecortin Family shRNA on Neuronal Migration Associated with Endogenous MicroRNA Dysregulation. Neuron 82, 1255–1262.

Bassett, A.R., Akhtar, A., Barlow, D.P., Bird, A.P., Brockdorff, N., Duboule, D., Ephrussi, A., Ferguson-Smith, A.C., Gingeras, T.R., Haerty, W., et al.(2014). Considerations when investigating lncRNA function in vivo. eLife *3*, e03058.

Bauer, M., Kinkl, N., Meixner, A., Kremmer, E., Riemenschneider, M., Forstl, H., Gasser, T., and Ueffing, M. (2009). Prevention of interferon-stimulated gene expression using microRNA-designed hairpins. Gene therapy 16, 142–147.

Belgard, T.G., Marques, A.C., Oliver, P.L., Abaan, H.O., Sirey, T.M., Hoerder-Suabedissen, A., Garcia-Moreno, F., Molnar, Z., Margulies, E.H., and Ponting, C.P. (2011). A Transcriptomic Atlas of Mouse Neocortical Layers. Neuron 71, 605–616.

Benavides-Piccione, R., Dierssen, M., Ballesteros-Yanez, I., Martinez de Lagran, M., Arbones, M.L., Fotaki, V., DeFelipe, J., and Elston, G.N. (2005). Alterations in the phenotype of neocortical pyramidal cells in the Dyrk1A+/-mouse. Neurobiology of disease 20, 115–122.

Berghoff, E.G., Clark, M.F., Chen, S., Cajigas, I., Leib, D.E., and Kohtz, J.D. (2013). Evf2 (Dlx6as) lncRNA regulates ultraconserved enhancer methylation and the differential transcriptional control of adjacent genes. Development 140, 4407–4416.

Boulard, M., Edwards, J.R., and Bestor, T.H. (2015). FBXL10 protects Polycomb-bound genes from hypermethylation. Nature genetics 47, 479–485.

Buchwald, G., van der Stoop, P., Weichenrieder, O., Perrakis, A., van Lohuizen, M., and Sixma, T.K. (2006). Structure and E3-ligase activity of the Ring-Ring complex of polycomb proteins Bmi1 and Ring1b. The EMBO journal 25, 2465–2474.

Cabianca, D.S., Casa, V., Bodega, B., Xynos, A., Ginelli, E., Tanaka, Y., and Gabellini, D. (2012). A long ncRNA links copy number variation to a polycomb/trithorax epigenetic switch in FSHD muscular dystrophy. Cell 149, 819–831.

Cheng, M., Jin, X., Mu, L., Wang, F., Li, W., Zhong, X., Liu, X., Shen, W., Liu, Y., and Zhou, Y. (2016). Combination of the clustered regularly interspaced short palindromic repeats (CRISPR)-associated 9 technique with the piggybac transposon system for mouse in utero electroporation to study cortical development. Journal of neuroscience research 94, 814–824.

Corish, P., and Tyler-Smith, C. (1999). Attenuation of green fluorescent protein half-life in mammalian cells. Protein Engineering, Design and Selection 12, 1035–1040.

Diez-Roux, G., Banfi, S., Sultan, M., Geffers, L., Anand, S., Rozado, D., Magen, A., Canidio, E., Pagani, M., Peluso, I., et al.(2011). A high-resolution anatomical atlas of the transcriptome in the mouse embryo. PLoS biology 9, e1000582.

Dillman, A.A., Hauser, D.N., Gibbs, J.R., Nalls, M.A., McCoy, M.K., Rudenko, I.N., Galter, D., and Cookson, M.R. (2013). mRNA expression, splicing and editing in the embryonic and adult mouse cerebral cortex. Nature neuroscience 16, 499–506.

Engreitz, J.M., Haines, J.E., Perez, E.M., Munson, G., Chen, J., Kane, M., McDonel, P.E., Guttman, M., and Lander, E.S. (2016). Local regulation of gene expression by lncRNA promoters, transcription and splicing. Nature 539, 452–455.

Farcas, A.M., Blackledge, N.P., Sudbery, I., Long, H.K., McGouran, J.F., Rose, N.R., Lee, S., Sims, D., Cerase, A., Sheahan, T.W., et al. (2012). KDM2B links the Polycomb Repressive Complex 1 (PRC1) to recognition of CpG islands. eLife 1, e00205.

Fietz, S.A., and Huttner, W.B. (2011). Cortical progenitor expansion, self-renewal and neurogenesis-a polarized perspective. Current opinion in neurobiology 21, 23–35.

Fotaki, V., Dierssen, M., Alcantara, S., Martinez, S., Marti, E., Casas, C., Visa, J., Soriano, E., Estivill, X., and Arbones, M.L. (2002). Dyrk1A haploinsufficiency affects viability and causes developmental delay and abnormal brain morphology in mice. Molecular and cellular biology 22, 6636–6647.

Fu, X.-D. (2014). Non-coding RNA: a new frontier in regulatory biology. National science review 1, 190–204.

Gaspard, N., Bouschet, T., Hourez, R., Dimidschstein, J., Naeije, G., van den Ameele, J., Espuny-Camacho, I., Herpoel, A., Passante, L., Schiffmann, S.N., et al.(2008). An intrinsic mechanism of corticogenesis from embryonic stem cells. Nature 455, 351–357.

Ghosh, A., and Greenberg, M.E. (1995). Distinct roles for bFGF and NT-3 in the regulation of cortical neurogenesis. Neuron 15, 89–103.

Goecks, J., Nekrutenko, A., Taylor, J., and Team, T.G. (2010). Galaxy: a comprehensive approach for supporting accessible, reproducible, and transparent computational research in the life sciences. Genome Biology 11, R86.

Greig, L.C., Woodworth, M.B., Galazo, M.J., Padmanabhan, H., and Macklis, J.D. (2013). Molecular logic of neocortical projection neuron specification, development and diversity. Nature Reviews Neuroscience 14, 755–769.

Grote, P., Wittler, L., Hendrix, D., Koch, F., Wahrisch, S., Beisaw, A., Macura, K., Blass, G., Kellis, M., Werber, M., et al.(2013). The tissue-specific lncRNA Fendrr is an essential regulator of heart and body wall development in the mouse. Developmental cell 24, 206–214.

Guo, C., Eckler, M.J., McKenna, W.L., McKinsey, G.L., Rubenstein, J.L., and Chen, B. (2013). Fezf2 expression identifies a multipotent progenitor for neocortical projection neurons, astrocytes, and oligodendrocytes. Neuron 80, 1167–1174.

Guttman, M., Garber, M., Levin, J.Z., Donaghey, J., Robinson, J., Adiconis, X., Fan, L., Koziol, M.J., Gnirke, A., Nusbaum, C., et al.(2010). Ab initio reconstruction of cell type-specific transcriptomes in mouse reveals the conserved multi-exonic structure of lincRNAs. Nat Biotech 28, 503–510.

Hagege, H., Klous, P., Braem, C., Splinter, E., Dekker, J., Cathala, G., de Laat, W., and Forne, T. (2007). Quantitative analysis of chromosome conformation capture assays (3C-qPCR). Nature protocols 2, 1722–1733.

He, J., Kallin, E.M., Tsukada, Y., and Zhang, Y. (2008). The H3K36 demethylase Jhdm1b/Kdm2b regulates cell proliferation and senescence through p15(Ink4b). Nature structural & molecular biology 15, 1169–1175.

He, J., Shen, L., Wan, M., Taranova, O., Wu, H., and Zhang, Y. (2013). Kdm2b maintains murine embryonic stem cell status by recruiting PRC1 complex to CpG islands of developmental genes. Nature cell biology 15, 373–384.

Heintzman, N.D., Hon, G.C., Hawkins, R.D., Kheradpour, P., Stark, A., Harp, L.F., Ye, Z., Lee, L.K., Stuart, R.K., and Ching, C.W. (2009). Histone modifications at human enhancers reflect global cell type-specific gene expression. Nature 459, 108.

Hirabayashi, Y., Suzki, N., Tsuboi, M., Endo, T.A., Toyoda, T., Shinga, J., Koseki, H., Vidal, M., and Gotoh, Y. (2009). Polycomb limits the neurogenic competence of neural precursor cells to promote astrogenic fate transition. Neuron 63, 600–613.

Homem, C.C., Repic, M., and Knoblich, J.A. (2015). Proliferation control in neural stem and progenitor cells. Nat Rev Neurosci 16, 647–659.

Huang, D.W., Sherman, B.T., and Lempicki, R.A. (2008). Systematic and integrative analysis of large gene lists using DAVID bioinformatics resources. Nat Protocols 4, 44–57.

Illingworth, R.S., Botting, C.H., Grimes, G.R., Bickmore, W.A., and Eskeland, R. (2012). PRC1 and PRC2 are not required for targeting of H2A.Z to developmental genes in embryonic stem cells. PloS one 7, e34848.

Illingworth, R.S., Moffat, M., Mann, A.R., Read, D., Hunter, C.J., Pradeepa, M.M., Adams, I. R., and Bickmore, W.A. (2015). The E3 ubiquitin ligase activity of RING1B is not essential for early mouse development. Genes Dev 29, 1897–1902.

Imayoshi, I., and Kageyama, R. (2014). bHLH factors in self-renewal, multipotency, and fate choice of neural progenitor cells. Neuron 82, 9–23.

Inagaki, T., Iwasaki, S., Matsumura, Y., Kawamura, T., Tanaka, T., Abe, Y., Yamasaki, A., Tsurutani, Y., Yoshida, A., Chikaoka, Y., et al.(2015). The FBXL10/KDM2B scaffolding protein associates with novel polycomb repressive complex-1 to regulate adipogenesis. The Journal of biological chemistry 290, 4163–4177.

Jiang, Y., Qian, X., Shen, J., Wang, Y., Li, X., Liu, R., Xia, Y., Chen, Q., Peng, G., Lin, S.Y., et al.(2015). Local generation of fumarate promotes DNA repair through inhibition of histone H3 demethylation. Nature cell biology 17, 1158–1168.

Kent, W.J., Sugnet, C.W., Furey, T.S., Roskin, K.M., Pringle, T.H., Zahler, A.M., and Haussler, D. (2002). The human genome browser at UCSC. Genome research 12, 996–1006.

Klattenhoff, C.A., Scheuermann, J.C., Surface, L.E., Bradley, R.K., Fields, P.A., Steinhauser, M.L., Ding, H., Butty, V.L., Torrey, L., Haas, S., et al.(2013). Braveheart, a long noncoding RNA required for cardiovascular lineage commitment. Cell 152, 570–583.

Kottakis, F., Foltopoulou, P., Sanidas, I., Keller, P., Wronski, A., Dake, B.T., Ezell, S.A., Shen, Z., Naber, S.P., Hinds, P.W., et al.(2014). NDY1/KDM2B functions as a master regulator of polycomb complexes and controls self-renewal of breast cancer stem cells. Cancer research 74, 3935–3946.

Kottakis, F., Polytarchou, C., Foltopoulou, P., Sanidas, I., Kampranis, S.C., and Tsichlis, P.N. (2011). FGF-2 regulates cell proliferation, migration, and angiogenesis through an NDY1/KDM2B-miR-101-EZH2 pathway. Molecular cell 43, 285–298.

Kwan, K.Y., Sestan, N., and Anton, E.S. (2012). Transcriptional co-regulation of neuronal migration and laminar identity in the neocortex. Development 139, 1535–1546.

Labonne, J.D., Lee, K.H., Iwase, S., Kong, I.K., Diamond, M.P., Layman, L.C., Kim, C.H., and Kim, H.G. (2016). An atypical 12q24.31 microdeletion implicates six genes including a histone demethylase KDM2B and a histone methyltransferase SETD1B in syndromic intellectual disability. Human genetics 135, 757–771.

Lepoivre, C., Belhocine, M., Bergon, A., Griffon, A., Yammine, M., Vanhille, L., Zacarias-Cabeza, J., Garibal, M.-A., Koch, F., Maqbool, M.A., et al.(2013). Divergent transcription is associated with promoters of transcriptional regulators. BMC Genomics 14, 914.

Li, H., Lai, P., Jia, J., Song, Y., Xia, Q., Huang, K., He, N., Ping, W., Chen, J., Yang, Z., et al. (2017a). RNA Helicase DDX5 Inhibits Reprogramming to Pluripotency by miRNA-Based Repression of RYBP and its PRC1-Dependent and -Independent Functions. Cell stem cell 20, 462–477. e466.

Li, W., Notani, D., Ma, Q., Tanasa, B., Nunez, E., Chen, A.Y., Merkurjev, D., Zhang, J., Ohgi, K., Song, X., et al.(2013). Functional roles of enhancer RNAs for oestrogen-dependent transcriptional activation. Nature 498, 516–520.

Li, Y., Wang, W., Wang, F., Wu, Q., Li, W., Zhong, X., Tian, K., Zeng, T., Gao, L., Liu, Y., et al. (2017b). Paired related homeobox 1 transactivates dopamine D2 receptor to maintain propagation and tumorigenicity of glioma-initiating cells. Journal of molecular cell biology.

Liang, G., He, J., and Zhang, Y. (2012). Kdm2b promotes induced pluripotent stem cell generation by facilitating gene activation early in reprogramming. Nature cell biology 14, 457–466.

Lin, M.F., Jungreis, I., and Kellis, M. (2011). PhyloCSF: a comparative genomics method to distinguish protein coding and non-coding regions. Bioinformatics (Oxford, England) 27, i275–282.

Lin, N.W., Chang, K.Y., Li, Z.H., Gates, K., Rana, Z.A., Dang, J.S., Zhang, D.H., Han, T.X., Yang, C.S., Cunningham, T.J., et al. (2014). An Evolutionarily Conserved Long Noncoding RNA TUNA Controls Pluripotency and Neural Lineage Commitment (vol 53, pg 1005, 2014). Molecular Cell 53, 1067–1067.

Liu, B., Ye, B., Yang, L., Zhu, X., Huang, G., Zhu, P., Du, Y., Wu, J., Qin, X., Chen, R., et al.(2017). Long noncoding RNA lncKdm2b is required for ILC3 maintenance by initiation of Zfp292 expression. Nature immunology 18, 499–508.

Luo, S., Lu, J.Y., Liu, L., Yin, Y., Chen, C., Han, X., Wu, B., Xu, R., Liu, W., Yan, P., et al.(2016). Divergent lncRNAs Regulate Gene Expression and Lineage Differentiation in Pluripotent Cells. Cell stem cell 18, 637–652.

Madisen, L., Zwingman, T.A., Sunkin, S.M., Oh, S.W., Zariwala, H.A., Gu, H., Ng, L.L., Palmiter, R.D., Hawrylycz, M.J., Jones, A.R., et al.(2010). A robust and high-throughput Cre reporting and characterization system for the whole mouse brain. Nature neuroscience 13, 133–140.

Mercer, T.R., Qureshi, I.A., Gokhan, S., Dinger, M.E., Li, G.Y., Mattick, J.S., and Mehler, M.F. (2010). Long noncoding RNAs in neuronal-glial fate specification and oligodendrocyte lineage maturation. Bmc Neuroscience 11.

Molyneaux, B.J., Goff, L.A., Brettler, A.C., Chen, H.H., Brown, J.R., Hrvatin, S., Rinn, J.L., and Arlotta, P. (2015). DeCoN: genome-wide analysis of in vivo transcriptional dynamics during pyramidal neuron fate selection in neocortex. Neuron 85, 275–288.

Morimoto-Suzki, N., Hirabayashi, Y., Tyssowski, K., Shinga, J., Vidal, M., Koseki, H., and Gotoh, Y. (2014). The polycomb component Ring1B regulates the timed termination of subcerebral projection neuron production during mouse neocortical development. Development 141, 4343–4353.

Ng, A., and Xavier, R.J. (2011). Leucine-rich repeat (LRR) proteins: integrators of pattern recognition and signaling in immunity. Autophagy 7, 1082–1084.

Ng, S.Y., Bogu, G.K., Soh, B.S., and Stanton, L.W. (2013). The Long Noncoding RNA RMST Interacts with SoX2 to Regulate Neurogenesis. Molecular Cell 51, 349–359.

Noonan, F.C., Goodfellow, P.J., Staloch, L.J., Mutch, D.G., and Simon, T.C. (2003). Antisense transcripts at the EMX2 locus in human and mouse. Genomics 81, 58–66.

ørom, U.A., Derrien, T., Beringer, M., Gumireddy, K., Gardini, A., Bussotti, G., Lai, F., Zytnicki, M., Notredame, C., Huang, Q., et al.(2010). Long Noncoding RNAs with Enhancer-like Function in Human Cells. Cell 143, 46–58.

Perino, M., and Veenstra, G.J. (2016). Chromatin Control of Developmental Dynamics and Plasticity. Dev Cell 38, 610–620.

Ponjavic, J., Oliver, P.L., Lunter, G., and Ponting, C.P. (2009). Genomic and transcriptional co-localization of protein-coding and long non-coding RNA pairs in the developing brain. PLoS genetics 5, e1000617.

Ramos, Alexander D., Diaz, A., Nellore, A., Delgado, Ryan N., Park, K.-Y., Gonzales-Roybal, G., Oldham, Michael C., Song, Jun S., and Lim, Daniel A. (2013). Integration of Genome-wide Approaches Identifies lncRNAs of Adult Neural Stem Cells and Their Progeny In Vivo. Cell stem cell 12, 616–628.

Rinn, J.L., and Chang, H.Y. (2012). Genome Regulation by Long Noncoding RNAs. Annual Review of Biochemistry, Vol 81 81, 145–166.

Roberts, T.C., Hart, J.R., Kaikkonen, M.U., Weinberg, M.S., Vogt, P.K., and Morris, K.V. (2015). Quantification of nascent transcription by bromouridine immunocapture nuclear run-on RT-qPCR. Nature protocols 10, 1198–1211.

Saba, L.M., Flink, S.C., Vanderlinden, L.A., Israel, Y., Tampier, L., Colombo, G., Kiianmaa, K., Bell, R.L., Printz, M.P., Flodman, P., et al.(2015). The sequenced rat brain transcriptome--its use in identifying networks predisposing alcohol consumption. The FEBS journal *282*, 3556–3578.

Scruggs, B.S., and Adelman, K. (2015). The Importance of Controlling Transcription Elongation at Coding and Noncoding RNA Loci. Cold Spring Harbor symposia on quantitative biology *80*, 33–44.

Scruggs, B.S., Gilchrist, D.A., Nechaev, S., Muse, G.W., Burkholder, A., Fargo, D.C., and Adelman, K. (2015). Bidirectional Transcription Arises from Two Distinct Hubs of Transcription Factor Binding and Active Chromatin. Molecular cell 58, 1101–1112.

Shen, Q., Wang, Y., Dimos, J.T., Fasano, C.A., Phoenix, T.N., Lemischka, I.R., Ivanova, N.B., Stifani, S., Morrisey, E.E., and Temple, S. (2006). The timing of cortical neurogenesis is encoded within lineages of individual progenitor cells. Nature Neuroscience 9, 743–751.

Sigova, A.A., Mullen, A.C., Molinie, B., Gupta, S., Orlando, D.A., Guenther, M.G., Almada, A. E., Lin, C., Sharp, P.A., Giallourakis, C.C., et al.(2013). Divergent transcription of long noncoding RNA/mRNA gene pairs in embryonic stem cells. Proceedings of the National Academy of Sciences of the United States of America 110, 2876–2881.

Sinnamon, J.R., Waddell, C.B., Nik, S., Chen, E.I., and Czaplinski, K. (2012). Hnrpab regulates neural development and neuron cell survival after glutamate stimulation. RNA (New York, NY) 18, 704–719.

Spigoni, G., Gedressi, C., and Mallamaci, A. (2010). Regulation of Emx2 expression by antisense transcripts in murine cortico-cerebral precursors. PloS one *5*, e8658.

Trapnell, C., Pachter, L., and Salzberg, S.L. (2009). TopHat: discovering splice junctions with RNA-Seq. Bioinformatics 25, 1105–1111.

Trapnell, C., Roberts, A., Goff, L., Pertea, G., Kim, D., Kelley, D.R., Pimentel, H., Salzberg, S.L., Rinn, J.L., and Pachter, L. (2012). Differential gene and transcript expression analysis of RNA-seq experiments with TopHat and Cufflinks. Nat Protocols 7, 562–578.

Trimarchi, T., Bilal, E., Ntziachristos, P., Fabbri, G., Dalla-Favera, R., Tsirigos, A., and Aifantis, I. (2014). Genome-wide Mapping and Characterization of Notch-Regulated Long Noncoding RNAs in Acute Leukemia Cell 158, 593–606.

Tsai, M.C., Manor, O., Wan, Y., Mosammaparast, N., Wang, J.K., Lan, F., Shi, Y., Segal, E., and Chang, H.Y. (2010). Long noncoding RNA as modular scaffold of histone modification complexes. Science 329, 689–693.

Venkov, C.D., Link, A.J., Jennings, J.L., Plieth, D., Inoue, T., Nagai, K., Xu, C., Dimitrova, Y.N., Rauscher, F.J., and Neilson, E.G. (2007). A proximal activator of transcription in epithelial-mesenchymal transition. The Journal of clinical investigation 117, 482–491.

Vescovi, A.L., Reynolds, B.A., Fraser, D.D., and Weiss, S. (1993). bFGF regulates the proliferative fate of unipotent (neuronal) and bipotent (neuronal/astroglial) EGF-generated CNS progenitor cells. Neuron 11, 951–966.

Vickers, T.A., Koo, S., Bennett, C.F., Crooke, S.T., Dean, N.M., and Baker, B.F. (2003). Efficient reduction of target RNAs by small interfering RNA and RNase H-dependent antisense agents A comparative analysis. Journal of Biological Chemistry 278, 7108–7118.

Vierstra, J., Rynes, E., Sandstrom, R., Zhang, M., Canfield, T., Hansen, R.S., Stehling-Sun, S., Sabo, P.J., Byron, R., Humbert, R., et al.(2014). Mouse regulatory DNA landscapes reveal global principles of *cis*-regulatory evolution. Science 346, 1007–1012.

Walder, R.Y., and Walder, J.A. (1988). Role of RNase H in hybrid-arrested translation by antisense oligonucleotides. Proceedings of the National Academy of Sciences 85, 5011–5015.

Wang, A., Wang, J., Liu, Y., and Zhou, Y. (2017). Mechanisms of Long Non-Coding RNAs in the Assembly and Plasticity of Neural Circuitry. Frontiers in neural circuits 11, 76.

Wang, K.C., Yang, Y.W., Liu, B., Sanyal, A., Corces-Zimmerman, R., Chen, Y., Lajoie, B. R., Protacio, A., Flynn, R.A., Gupta, R.A., et al. (2011a). A long noncoding RNA maintains active chromatin to coordinate homeotic gene expression. Nature 472, 120–U158.

Wang, L., Park, H.J., Dasari, S., Wang, S., Kocher, J.P., and Li, W. (2013). CPAT: Coding-Potential Assessment Tool using an alignment-free logistic regression model. Nucleic acids research 41, e74.

Wang, T., Chen, K., Zeng, X., Yang, J., Wu, Y., Shi, X., Qin, B., Zeng, L., Esteban, M.A., Pan, G., et al. (2011b). The histone demethylases Jhdm1a/1b enhance somatic cell reprogramming in a vitamin-C-dependent manner. Cell stem cell 9, 575–587.

Wang, Y., He, L., Du, Y., Zhu, P., Huang, G., Luo, J., Yan, X., Ye, B., Li, C., Xia, P., et al.(2015). The Long Noncoding RNA lncTCF7 Promotes Self-Renewal of Human Liver Cancer Stem Cells through Activation of Wnt Signaling. Cell stem cell 16, 413–425.

Wu, M., Wang, P.F., Lee, J.S., Martin-Brown, S., Florens, L., Washburn, M., and Shilatifard, A. (2008). Molecular regulation of H3K4 trimethylation by Wdr82, a component of human Set1/COMPASS. Molecular and cellular biology 28, 7337–7344.

Wu, X., Johansen, J.V., and Helin, K. (2013). Fbxl10/Kdm2b Recruits Polycomb Repressive Complex 1 to CpG Islands and Regulates H2A Ubiquitylation. Molecular cell.

Xing, Y.H., Yao, R.W., Zhang, Y., Guo, C.J., Jiang, S., Xu, G., Dong, R., Yang, L., and Chen, L.L. (2017). SLERT Regulates DDX21 Rings Associated with Pol I Transcription. Cell 169, 664–678.e616.

Ye, B., Liu, B., Yang, L., Zhu, X., Zhang, D., Wu, W., Zhu, P., Wang, Y., Wang, S., Xia, P., et al.(2018). LncKdm2b controls self-renewal of embryonic stem cells via activating expression of transcription factor Zbtb3. The EMBO journal *37*.

Yin, Y., Yan, P., Lu, J., Song, G., Zhu, Y., Li, Z., Zhao, Y., Shen, B., Huang, X., Zhu, H., et al.(2015). Opposing Roles for the lncRNA Haunt and Its Genomic Locus in Regulating HOXA Gene Activation during Embryonic Stem Cell Differentiation. Cell stem cell 16, 504-516.

Yue, F., Cheng, Y., Breschi, A., Vierstra, J., Wu, W., Ryba, T., Sandstrom, R., Ma, Z., Davis, C., Pope, B.D., et al.(2014). A comparative encyclopedia of DNA elements in the mouse genome. Nature 515, 355–364.

Zhou, Z.-J., Dai, Z., Zhou, S.-L., Hu, Z.-Q., Chen, Q., Zhao, Y.-M., Shi, Y.-H., Gao, Q., Wu, W.-Z., Qiu, S.-J., et al.(2014). HNRNPAB Induces Epithelial–Mesenchymal Transition and Promotes Metastasis of Hepatocellular Carcinoma by Transcriptionally Activating SNAIL. Cancer research 74, 2750–2762.

